# Post-translational acylation drives folding and activity of the CyaA bacterial toxin

**DOI:** 10.1101/2025.09.11.673628

**Authors:** Corentin Léger, Samuel E. Hoff, Gaia Scilironi, Amiel Abettan, Roshan Shrestha, Jacinthe Frangieh, Vincent Deruelle, Nicolas Carvalho, Dorothée Raoux-Barbot, Nathalie Duclert-Savatier, Benjamin Bardiaux, Sébastien Brier, François Bontems, Gérard Pehau-Arnaudet, Daniel Ladant, Luca Monticelli, Massimiliano Bonomi, Alexandre Chenal

## Abstract

Post-translational modifications critically shape protein conformation and function, yet how they regulate bacterial toxins remains elusive. The adenylate cyclase (CyaA) toxin is a major virulence factor of *Bordetella pertussis*, the causative agent of whooping cough. CyaA is produced as an inactive precursor, proCyaA, which is activated by acylation of two lysine residues within the bacterium. Once acylated and secreted, CyaA invades innate immune cells and disrupts their phagocytic functions. High-resolution structural characterization of CyaA has remained elusive due to its size, multi-domain organization, flexibility, and aggregation propensity. Here, we overcome these challenges and generate the first structural ensembles of both non-acylated and acylated CyaA in solution by combining experimental data with integrative modeling. Coarse-grained molecular dynamics simulations reveal that acylation is critical to stabilize the native fold and to favorably orient CyaA on the target membrane. Overall, our findings reveal how post-translational acylation triggers native folding and provide mechanistic insights into the early steps of host cell intoxication.

## Introduction

The adenylate cyclase toxin (CyaA) is a key virulence factor of *Bordetella Pertussis*, the bacterium that causes whooping cough, a highly contagious disease that can have dramatic consequences for vulnerable population such as children^1^. Despite global vaccination efforts, whooping cough remains a significant public health concern, causing an estimated 160,000 deaths annually among children under the age of five, with the vast majority occurring in low- and middle-income countries^2^. Infants under one year are particularly at risk, accounting for most pertussis-related fatalities even in high-income countries. CyaA plays an essential role in the early stage of the respiratory tract infection by *B. pertussis*: it invades innate immune cells (e.g., alveolar macrophages and neutrophils), where it catalyzes a massive overproduction of cAMP. This disrupts cellular function, paralyzing the immune response and eventually leading to cell death^3^.

CyaA is a 1706-residue protein belonging to the Repeat-in-ToXin (RTX) cytolysin family^4^. These virulence factors, produced by many Gram-negative pathogens, form cation-selective pores in the plasma membranes of specific target cells. The RTX toxins share several common features^5^. First, their C-terminal domain is made of tandem repetitions of nonapeptide RTX motifs that are involved in calcium binding. The CyaA RTX domain (RD) contains ∼40 copies of such RTX motifs (residues 913-1613), which are disordered in the low-calcium environment of the bacterial cytosol^6,7^. After toxin secretion *via* a type I secretion machinery, they can fold in the extracellular medium into β-roll structures stabilized by calcium binding^7–9^. Second, the RTX cytolysins are post-translationally acylated on one or two specific lysine residues, K860 and K983 in the case of CyaA, located in the so-called acylation region (AR) of CyaA^10,11^. This acylation, carried out by a dedicated acyltransferase prior to secretion^12,4,5^, is essential for toxin biological activities^12^. Finally, RTX toxins contain a hydrophobic region (HR) composed of several hydrophobic and amphiphilic α-helices that can insert into the plasma membrane of target cells to form cation-selective pores^13^. In addition to these common characteristics, CyaA has an N-terminal catalytic domain (ACD) that displays a potent calmodulin (CaM)-dependent adenylate cyclase activity^14,15^, as well as a translocating region (TR)^16,17^, critical for ACD entry into target cells^18,19^.

CyaA intoxicates cells through a mechanism that is unique among all known bacterial toxins (**Fig. 1**): it is independent of receptor-mediated endocytosis and instead involves a direct translocation of its ACD across the plasma membrane of target cells^20^. In the currently established model, CyaA first binds in a calcium-dependent manner to the CD11b/CD18 integrin at the surface of innate immune cells *via* RD. It then inserts its hydrophobic and amphiphilic segments (TR, HR, and AR) into the plasma membrane of the target cell and subsequently translocate the ACD domain directly across the membrane into the cytosol^21^. There, ACD is refolded and activated upon binding to CaM^22^, leading to the production of supraphysiological levels of cAMP. Although this model outlines the overall sequence of events, the detailed molecular mechanism of CyaA entry remains largely unknown. In particular, while acylation of CyaA is essential for cellular intoxication^23^, the precise role of this post-translational modification in the cell invasion process is still unclear^12,11,24,20^.

**Figure 1.**
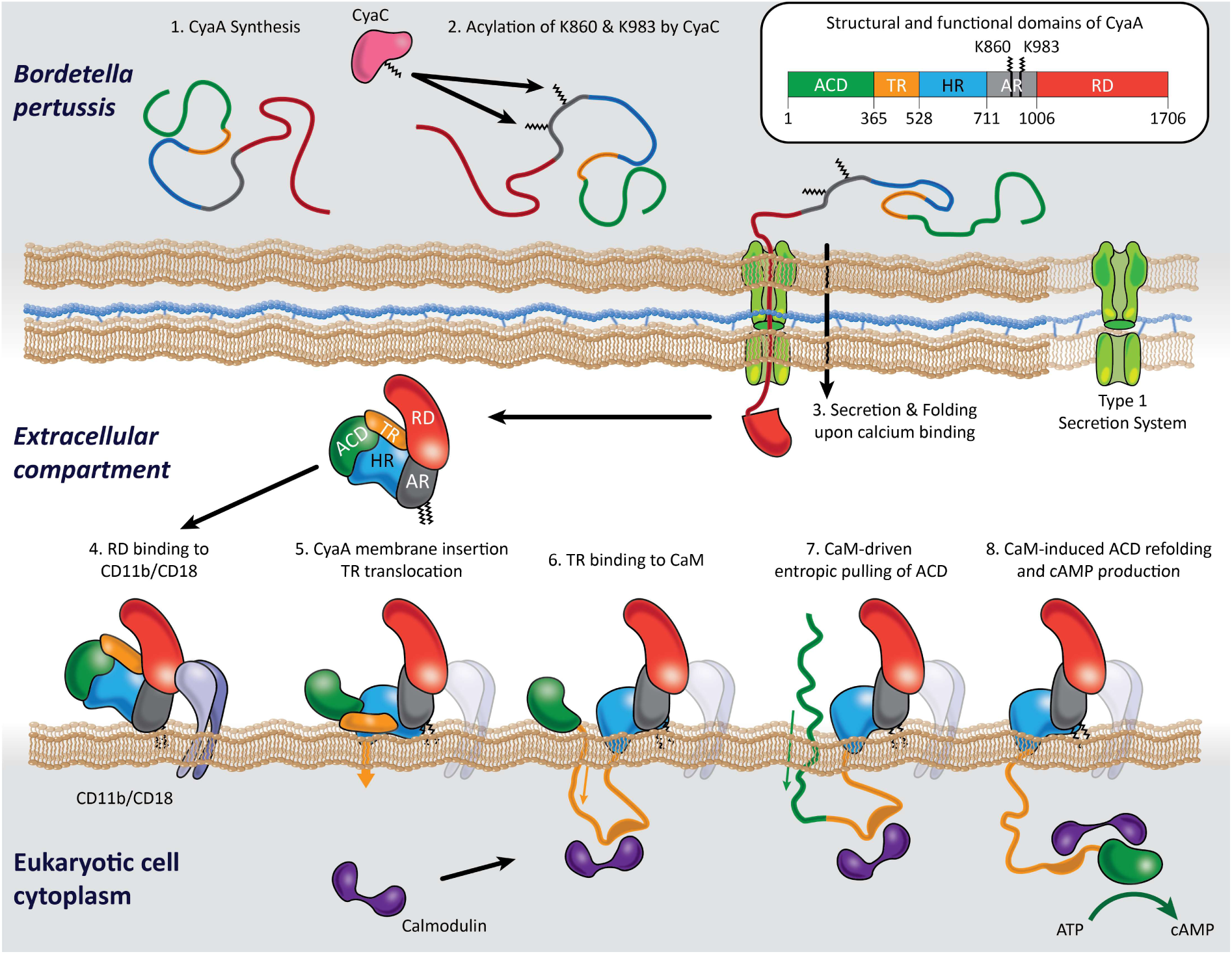
Model of the eucaryotic cell intoxication process by the CyaA toxin. The CyaA toxin is produced by *Bordetella pertussis* (1). CyaA is composed of five structural and functional domains (inset): the adenylate cyclase domain (ACD, green), the translocation region (TR, orange), the hydrophobic region (HR, cyan), the acylation region (AR, grey), and the cell-receptor binding domain (RD, red). The proCyaA protoxin is acylated by CyaC on lysine residues K860 and K983 (2). The CyaA toxin is then secreted and folded upon calcium binding in the extracellular compartment (3). The folded, calcium-loaded and acylated CyaA binds to eukaryotic cells (4) and translocate TR and ACD across the plasma membrane (5-7). Calmodulin-binding to ACD induces the refolding and activation of the enzyme, leading to the production of supraphysiological cAMP levels and cell death (8).

Although the role of CyaA in the infection and disease pathogenesis is widely documented^25,26^, high-resolution structural characterization of the full-length protein has remained elusive due to its large size, multi-domain organization, flexibility, and aggregation propensity. Yet, structural information is available only for fragments or domains of CyaA. Notably, Tang and colleagues solved the X-ray structure of ACD in complex with calmodulin^27^, while, more recently, Goldsmith *et al.* reported the Cryo-EM structure of the C-terminus RTX domain in complex with the extracellular domains of CD11b/CD18^28^. Additionally, in-solution studies have highlighted the structural dynamics and ligand-induced folding of ACD, TR and RD^7,8,22,29^.

Here, we build on our recent achievement in refolding CyaA into a stable, monomeric and functional form^30,31^ to characterize both the non-acylated (proCyaA) and acylated CyaA proteins using a complementary set of experimental techniques, including Small-Angle X-ray Scattering (SAXS), Hydrogen-Deuterium Exchange Mass Spectrometry (HDX-MS), and Cryo- Electron Microscopy (Cryo-EM). We develop an integrative approach that leverages the potential of raw single-particle Cryo-EM images to capture the full conformational landscape of dynamic proteins, enabling the combination of diverse experimental datasets to generate the first structural ensembles of CyaA and proCyaA in solution (**Extended Data** Fig. 1). These models allow us to identify the hydrophobic pockets accommodating the two acyl chains and to perform coarse-grained Molecular Dynamics (MD) simulations of both proteins in complex with the CD11b/CD18 integrin embedded in a lipid bilayer. This analysis reveals a critical role for the acyl chains, not only in stabilizing the native, functional fold of CyaA, but also in properly orienting the toxin to the membrane following receptor binding. Overall, this study provides key insights into how acylation governs the folding and biological activity of CyaA, with broader implications for understanding other RTX toxins and post-translationally acylated proteins.

## Results

### Production, purification, and biophysical characterization of proCyaA and CyaA

The recombinant proteins proCyaA (non-acylated) and CyaA (acylated on K860 and K983 upon co-expression of CyaC) were expressed in *Escherichia coli* and purified from inclusion bodies after solubilization in 6M Urea^30^. The purified proteins (**Extended Data** Fig. 2**.A**) were finally refolded in the presence of calcium via a molecular confinement procedure^30^ into monomeric forms that remain stable and monodisperse in physiological buffer without chaotropic agents, as shown by SEC analysis^31^ and Mass Photometry (**Extended Data** Fig. 2**.B, C**). A sensitive GloSensor™ cAMP assay^32^ revealed that the monomeric CyaA triggered massive accumulation of cAMP in intoxicated cells, whereas the monomeric proCyaA had no effect (**Extended Data** Fig. 2**.D, E**), confirming the functionality of the monomeric CyaA and the lack of invasion capacity of the non-acylated proCyaA. The structural properties of both monomeric proteins in solution were initially characterized by SAXS. Parameters inferred from SEC-SAXS analysis indicate that proCyaA (SASBDB accession code: SASDXH6) and CyaA (SASBDB accession code: SASDXG6) adopt similar shapes at the resolution level provided by SAXS analysis with R_g_/D_max_ of 48/175Å and 47/165Å, respectively (**Extended Data** Fig. 3 and **Supplementary Table 1**) and are similar to our previously published SEC-SAXS data on CyaA^31^.

### Cryo-EM analysis reveals conformational heterogeneity in CyaA and proCyaA

All attempts to obtain crystals suitable for X-ray crystallography from the CyaA or proCyaA monomers were unsuccessful, likely due to the flexibility and structural dynamics of these proteins. We therefore turned to Cryo-EM to obtain structural information. Cryo-EM images of CyaA and proCyaA were acquired on a Glacios Cryo-TEM (**Fig. 2**) and show monodisperse single particles with dimensions around 150-200Å. Picked particles were subjected to several rounds of reference free 2D classifications (**Fig. 2.A**). The 2D class averages show an ensemble of different conformations with well-defined portions and blurry edges. Due to this conformational heterogeneity, the final ensembles of particles for CyaA and proCyaA resulted in refined densities with a medium resolution of 6-8Å (**Fig. 2.B, C**). Because of the low resolution, the 3D reconstructions of CyaA and proCyaA did not offer more detailed structural information than the molecular envelopes obtained from SEC-SAXS data (**Extended Data** Fig. 3**.G, H**).

**Figure 2.**
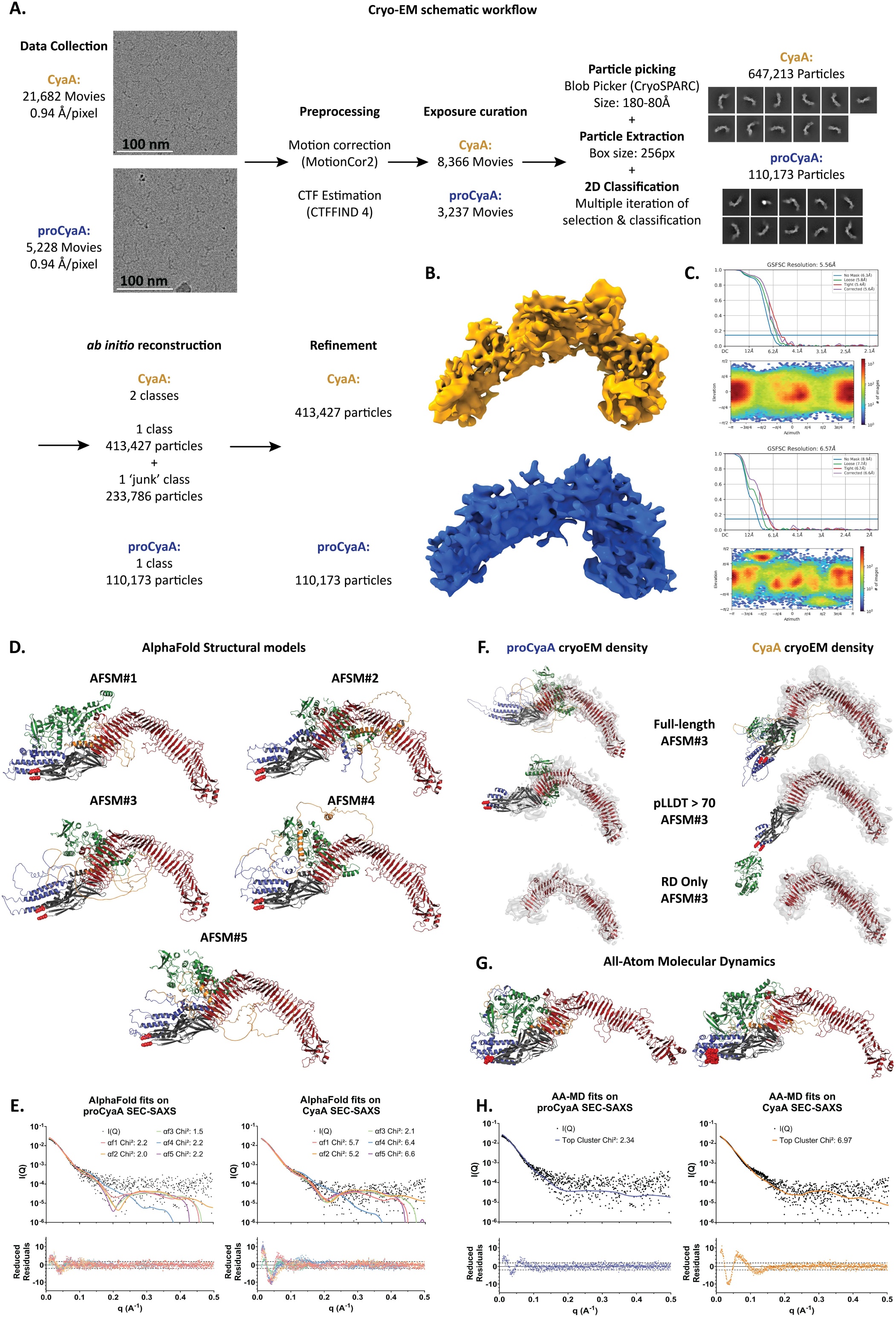
Cryo-EM data collection and initial structural modeling of proCyaA and CyaA. A. Schematic workflow of the cryo-EM single-particle analysis, resulting in one major class composed of 110,173 and 413,127 images for proCyaA and CyaA, respectively. B. Final density map of CyaA (top, yellow) and proCyaA (bottom, blue) at 5.6 and 6.6 Å resolution, respectively. C. Gold-Standard Fourier Shell Correlation (top) and Angular distribution of projection (bottom). Average 2D Classification and Angular Distribution of projection show a lack of particles in some orientation, indicating preferential orientations of CyaA and proCyaA on the grids. D. Top5 AlphaFold models of proCyaA in cartoon representation with ACD in green, TR in orange, HR in blue, AR in black, and RD in dark red with lysine residues K860 and K983 represented as red spheres. E. Theoretical SAXS profiles computed from AlphaFold models and comparison with experimental curves (top) along with reduced residuals (bottom) for proCyaA (left) and CyaA (right). Theoretical and experimental SAXS profiles are represented by black dots and colored lines, respectively. F. AlphaFold structural model #3 (AFSM#3) flexible fit inside the cryo-EM density map of proCyaA (left) and CyaA (right). Same color code as in panel D. G. All Atom- Molecular Dynamics (AA-MD) top cluster of proCyaA (left) and CyaA (right). Same color code as in panel D. H. Theoretical SAXS profiles computed from the AA-MD models and comparison to experimental curves of proCyaA (left) and CyaA (right), represented as in panel E.

Noteworthy, the molecular volumes computed by the electronic densities were much smaller than those computed from the SAXS data. The Cryo-EM 3D reconstructions match mainly RD that is known to be tightly folded in the presence of calcium^7^. The missing density and the low-resolution of CyaA and proCyaA proteins are likely caused by the structural dynamics of the other domains, as previously shown for ACD^22^ and TR^29^, resulting in high conformational heterogeneity. Additionally, preferential orientations of proteins on the grids, which limited certain views as seen in the 2D classes, may further reduce the overall resolution (**Fig. 2.A**).

### Limitations of AlphaFold models highlight the need for integrative ensemble refinement

Initial structural modeling of CyaA was performed using AlphaFold 2.2 (AF)^33^. The five best AF models of CyaA were structurally similar and comprised four well-folded domains (ACD, HR, AR and RD) along with one predominantly disordered domain (TR) (**Fig 2.D**). Since AlphaFold cannot model post-translational acylation on lysine residues, the resulting models likely represent the non-acylated proCyaA protein more accurately than the acylated CyaA. Nevertheless, these AlphaFold models provide a solid starting point for subsequent structural refinement of both CyaA and proCyaA.

We attempted to refine the AF models of proCyaA and CyaA using the SEC-SAXS curves and the cryo-EM density, as follows. PEPSI-SAXS was first used to generate theoretical SAXS scattering curves for all five AF models (**Fig 2.E**) with mean chi² values of 2.0 ± 0.3 and 5.2 ± 1.8 for proCyaA and CyaA, respectively. We then selected the AF model (AFSM#3) with best agreement with SEC-SAXS and previously published AUC and SEC-TDA for both CyaA and proCyaA datasets^30,31,24^ (**Supplementary Table 1 and 2**). Finally, iMODfit was used to flexibly refine AFSM#3 inside the cryo-EM density of CyaA and proCyaA. Although various fitting attempts were made using different segments of AFSM#3, selected based on AlphaFold confidence scores, none yielded satisfactory results. Overall, AFSM#3 could not be accommodated within the cryo-EM density of either the CyaA or proCyaA proteins (**Fig. 2.F**).

Since AF2 could not predict the folding of CyaA with acyl chains, acylations (C16:1 Δ9) were added *a posteriori* to AFSM#3 on both residues K860 and K983, along with the calcium ions that bind to each RTX motifs in RD to stabilize the beta-roll fold. Unrestrained all-atom molecular dynamics (AA MD) simulations in explicit solvent were then performed to assess whether the acylated AFSM#3 could adopt a more compact fold, yielding conformations more consistent with the low-resolution structures derived from SEC-SAXS and cryo-EM data. However, despite improving the fit to the SEC-SAXS curves, the structures sampled in AA MD simulations remained more extended than indicated by SEC-SAXS and cryo-EM data of CyaA (**Fig. 2.G,H**), prompting us to adopt a different integrative modeling approach, as detailed in the next section.

### Integrative structural ensembles of CyaA and proCyaA

So far, we have not succeeded in obtaining high-resolution cryo-EM structures of CyaA and proCyaA, nor to refine AlphaFold models using SAXS data and low-resolution cryo-EM maps. We have already emphasized that this task is particularly difficult for proteins that are highly dynamic and composed of multiple domains. To overcome these challenges, we developed an integrative modeling approach that combines coarse-grained physico-chemical models, HDX-MS data, and the raw single-particle cryo-EM images. This approach aims to capture the complete range of conformations that these dynamic proteins adopt. At the core of our method is BioEM^34^, a Bayesian refinement tool that begins with a set of initial structural models and calculates the likelihood of observing the raw cryo-EM images based on these models. This process ultimately enables us to estimate the relative population of each structural model within the sample, providing an accurate description of both the structural diversity and thermodynamic properties of dynamic ensembles.

We generated an initial ensemble using Martini coarse-grained simulations, starting from the AlphaFold models AFSM#3 we had previously obtained. Because Martini employs an elastic network to maintain the protein’s secondary structure, it inherently limits the sampling of the full conformational landscape. To address this issue, we used our previously published HDX- MS data^24^ to selectively remove elastic restraints on the highly dynamic segments of CyaA and proCyaA (Methods). From these coarse-grained simulations, we then clustered the conformations of CyaA and proCyaA, yielding a limited set of representative structures that capture the broad conformational landscapes of these two proteins.

We then used BioEM to estimate the likelihood of each individual cryo-EM single-particle image given each representative model, and subsequently to determine the weighted population of all CyaA and proCyaA models (Methods). In the following, we refer to our entire pipeline as Hc2B, which stands for refining **<u>H</u>**DX-MS relaxed **<u>c</u>**oarse-grained MD simulations with **<u>2</u>**D cryo-EM particle images using **<u>B</u>**ioEM. Overall, the Hc2B pipeline generated 121 models for CyaA and 127 models for proCyaA (**Fig. 3.A**). These models included compact as well as more extended conformations, with radius of gyration (R_g_) estimated by CRYSOL (Methods) covering the range 44-53Å and 39-50Å for CyaA and proCyaA, respectively (**Supplementary** Fig 2). Similarly, the largest interatomic distance (D_max_) spanned the range 162-231Å and 133- 202 Å for CyaA and proCyaA, respectively (**Supplementary** Fig 2). In each ensemble, one model was identified by BioEM to have a dominant population: model 4 (proCyaA_Hc2B_, population 52%, SASBDB accession code: SASDXH6) for proCyaA and model 100 (CyaA_Hc2B_, population 68%, SASBDB accession code: SASDXG6) for CyaA (**Fig. 3.B**).

**Figure 3.**
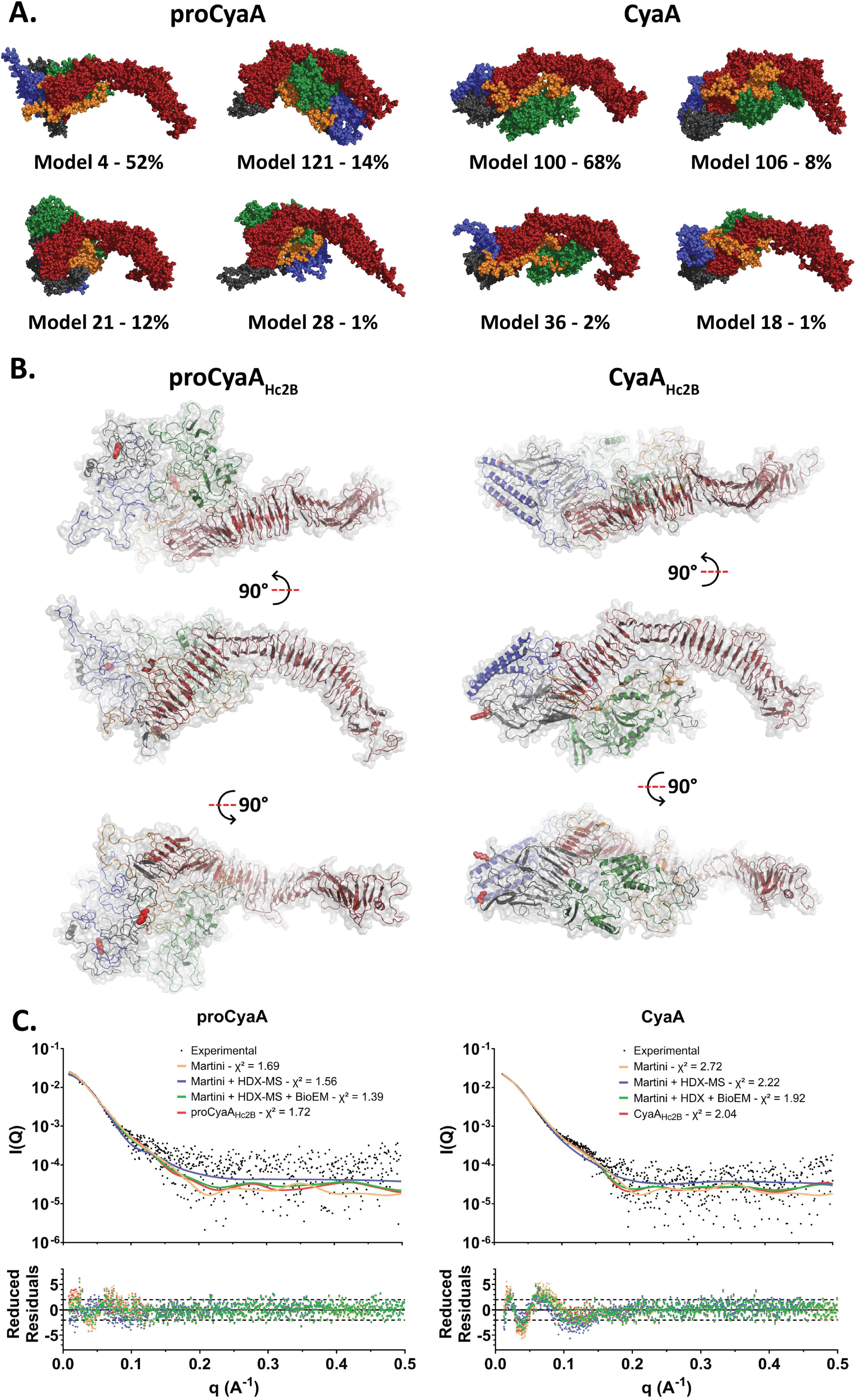
Integrative structural ensemble determination of proCyaA and CyaA. A. Four most represented models for proCyaA (left) and CyaA (right) with their associated population as determined from the cryo-EM single-particle images. Models are visualized using a surface representation with ACD in green, TR in orange, HR in blue, AR in black, and RD in dark red, with lysine residues K860 and K983 represented as red spheres. B. Most populated models obtained from the Hc2B integrative approach for proCyaA_Hc2B_ (left, SASBDB accession code: SASDXH6) and CyaA_Hc2B_ (right, SASBDB accession code: SASDXG6). All models are represented as in panel A. C. Comparison between experimental SAXS curves and theoretical profiles computed from different structural ensembles of proCyaA (left) and CyaA (right): standard Martini simulations (orange), HDX-MS Martini simulations (purple), and HDX-MS Martini ensemble refined by cryo-EM single-particles images (Hc2B, green). In red, theoretical SAXS curves of proCyaA_Hc2B_ and CyaA_Hc2B_, the most populated models as determined by our Hc2B integrative approach. Experimental SAXS curve is represented by black dots.

To validate the structural ensembles of CyaA and proCyaA generated by our integrative Hc2B approach, we compared the experimental SAXS and hydrodynamic data to theoretical SAXS curves computed from the full set of models produced by our pipeline, weighted according to the thermodynamic populations estimated by BioEM (Methods). As shown in **Fig. 3.C** and **Supplementary Table 1 and 2**, the Hc2B ensembles agree more closely with the experimental SAXS profile than the initial AFSM#3 models, the structural ensembles derived from standard Martini simulations, and the HDX-MS–relaxed coarse-grained models that were not reweighted using BioEM. Notably, standard Martini simulations without the HDX-MS–guided tuning of the elastic network show the poorest agreement with SAXS (**Supplementary Table 2**). Overall, this validation demonstrates that starting from a purely *in silico* ensemble, the stepwise integration of experimental data - first HDX-MS, then raw single-particle cryo-EM images - progressively improves the quality of the structural ensembles, as reflected by the increasingly better agreement with independent experimental data not used in the modeling process.

### Integrative structural ensembles reveal acylation-driven conformational reorganization in CyaA

The differences between our integrative proCyaA and CyaA ensembles illustrate the substantial impact of post-translational acylations on protein folding (**Fig. 3.B & 4.A,B**). The RD domain in CyaA_Hc2B_ and proCyaA_Hc2B_ displays a similar organization, with five well-defined RTX blocks that closely resemble those described by Goldsmith *et al*^28^. In CyaA_Hc2B_, the AR domain folds into a beta-rich structure that caps the N-terminal end of RD (**Fig. 4.A**), similar to the organization reported by Goldsmith *et al*. However, it differs notably at the tip near the acylation site, most likely because their AR-RD CyaA fragment was not acylated^28^. Interestingly, the two acylated lysine residues, K860 and K983, lie within 2 nm of the tips of the hydrophobic helices in HR. This local arrangement in CyaA_Hc2B_ creates a hydrophobic environment that may favor the folding of the surrounding apolar segments and promote membrane interaction. Most importantly, this tertiary structure organization and the associated interdomain interactions are absent in proCyaA_Hc2B_ (**Fig. 3.B**). In particular, HR is well folded in CyaA_Hc2B_, forming two long helices, but remains poorly structured in proCyaA_Hc2B_ (**Fig. 3.B & 4.A**). Finally, ACD is positioned in the groove formed by AR and RD in CyaA_Hc2B_ (**Fig. 3.B**), while the translocation region (TR) appears as an extended and disordered linker between ACD and HR.

**Figure 4:**
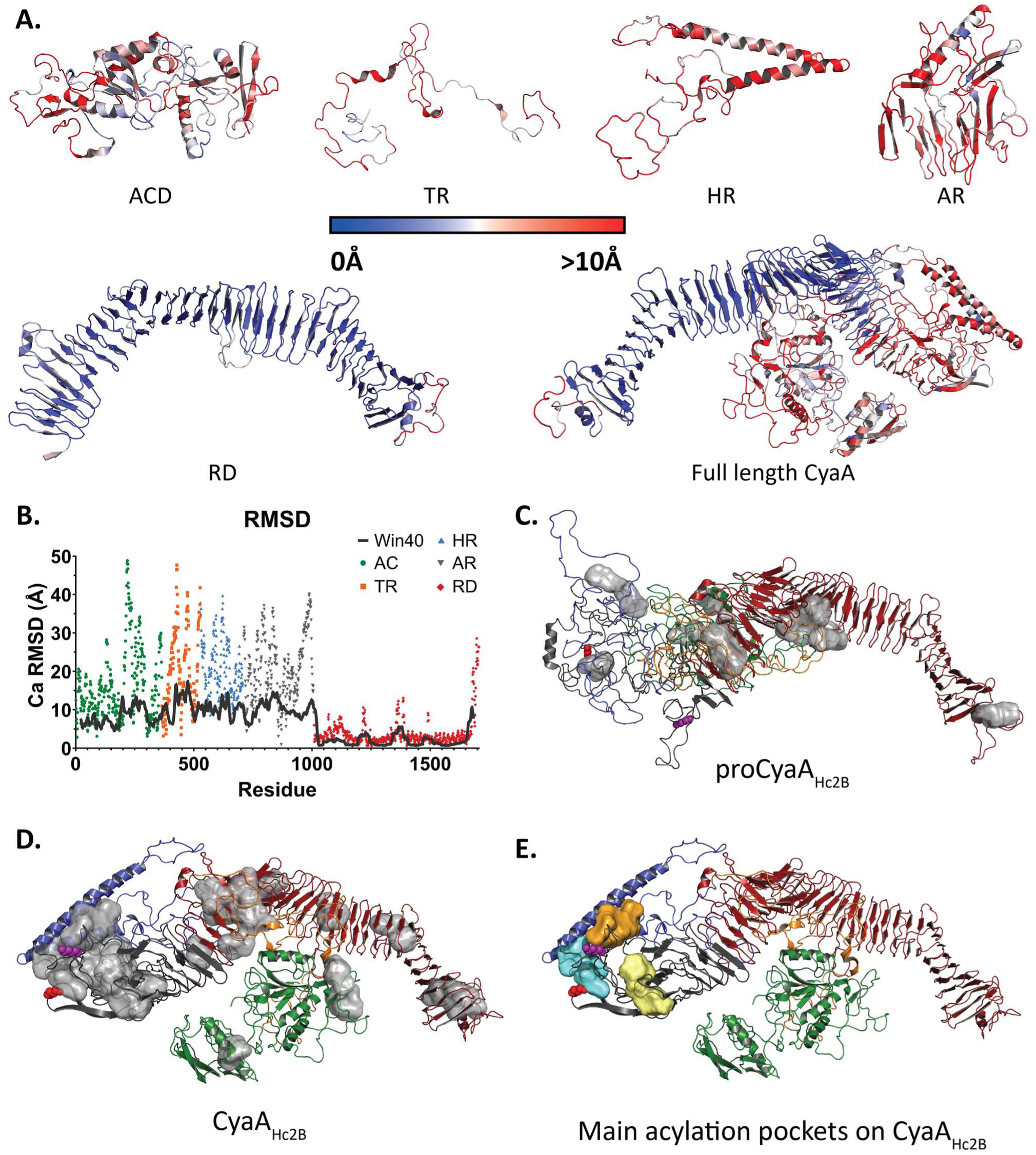
Analysis of the Hc2B models of proCyaA and CyaA. A. Similarity between proCyaA_Hc2B_ and CyaA_Hc2B_ models as measured by their C⍺ Root Mean Square Deviation (RMSD). Models are colored from blue to white to red based on the RMSD in a range between 0 to >10 Å. Individual domains as well as the full-length CyaA_Hc2B_ model are shown. B. Per-residue and 40-residue rolling mean RMSD between proCyaA_Hc2B_ and CyaA_Hc2B_ models across each domain. C & D. Cavities on proCyaA_Hc2B_ (C) and CyaA_Hc2B_ (D) with volumes large enough to accommodate the acylations. All identified pockets are colored in grey. E. Main acylation pockets identified by comparing CyaA_Hc2B_ with previously published HDX-MS data (see Extended Figure 4 and 5). Proteins are visualized with a cartoon representation with ACD in green, TR in orange, HR in blue, AR in black, and RD in dark red. Pockets P1, P2 and P3 are colored in blue, yellow and orange, respectively. The proCyaA and CyaA data are available on SASBDB with the following accession code: SASDXH6 and SASDXG6, respectively.

To quantitatively highlight the impact of acylation on the structures of CyaA and proCyaA, we computed the RMSD between each domain of CyaA_Hc2B_ and proCyaA_Hc2B_ and visualized the differences using color-coding in **Fig. 4.A**. These results provide direct evidence that RD is largely unaffected by acylation, while the structures of the other domains are all significantly altered (**Fig. 4.A, B**). We therefore concluded that acylation at K860 and K983 induces long-range allosteric effects on ACD, TR, and HR domains, far from the modification sites. Finally, we identified residues in proCyaA_Hc2B_ and CyaA_Hc2B_ (**Fig. 4.C, D**) that contribute to the formation of potential hydrophobic pockets capable of accommodating the acyl chains (C16:1 Δ9 in our model) attached to K860 and K983 of CyaA_Hc2B_. Three main pockets were mapped on CyaA_Hc2B_ (**Fig. 4.E**): the K860 acyl chain (red) fits well into the pocket 1 (P1, blue), while the K983 acyl chain (purple) could be accommodated by either pocket 2 (P2, yellow) or 3 (P3, orange) pockets. To probe this further, in the next section we used HDX-MS to identify acyl chain binding sites and their contribution to CyaA folding.

### HDX-MS identifies acyl chain binding sites and their role in CyaA folding

To further localize the hydrophobic pockets accommodating the K860 and K983 acyl chains and to decipher the contribution of acylations on CyaA refolding, we re-analyzed previously published HDX-MS data on CyaA and proCyaA^24^. In that work, HDX-MS data was used to probe the structural dynamics of both proCyaA and CyaA at various urea concentration. This study revealed dramatic differences in deuterium uptake between CyaA and proCyaA, primarily located in the hydrophobic and acylation region. Here, we mapped the HDX-MS uptake differences between CyaA and proCyaA at different urea concentration on our integrative CyaA_Hc2B_ model (**Extended Data** Fig. 4). As illustrated in **Extended Data** Fig. 5, the acyl chains on K860 and K983 residues notably stabilized the hydrophobic helical hairpin of HR in CyaA, as well as the segments surrounding the P1, P2, and P3 pockets in the AR domain.

Consistent with our integrative models of CyaA, the HDX-MS data allow us to propose that pocket P1 is a strong candidate to accommodate the acyl chain of K860, while pockets P2 and P3 may alternatively and dynamically bind the acyl chain of K983. This interaction appears critical for stabilizing the AR domain and maintaining the native folded state of CyaA (**Fig. 4.D**). Moreover, these data help explain why acylation at K983 is essential to achieve a functional toxin. While the absence of K860 acylation does not prevent AR refolding, which happens at lower urea concentration in proCyaA than in CyaA, the lack of K983 acylation prevents proCyaA from adopting the native fold observed in fully acylated CyaA in the absence of urea (**Extended Data** Fig. 5).

### Molecular Dynamics simulations reveal acylation-mediated stabilization and membrane anchoring of CyaA

To investigate the effects of acylation on the structural dynamics of the toxin, we linked acyl chains to K860 and K983 in the CyaA_Hc2B_ model (yielding CyaA_Hc2B_^acyl^) and performed coarse-grained molecular dynamics simulations of both proCyaA_Hc2B_ and CyaA_Hc2B_^acyl^ in solution (**Fig. 5.A** and **Supplementary Movies 1 and 2**). Interestingly, during MD simulations, while the acylated K860 resides mostly inside the P1 pocket (about 65% of the time), the acylated K983 appears to switch dynamically from P2 to P3 (yellow and orange, respectively in **Fig. 5.A**) pockets, with a preference for P3 (about 60% of the time) and some simultaneous occupation of P2 and P3 (about 3% of the time), due to partial overlap of both pockets (**Supplementary Movie 3** and **Supplementary Table 4**). The two acyl chains also make transient contact with one another, about 20% of the simulation time, and remain very dynamic. Root-mean square fluctuation (RMSF) analysis (**Fig. 5.B**) reveals significant differences in the flexibility of the TR, HR, and AR (residues 500-1000) domains (**Fig. 5.B**), with increased flexibility in proCyaA compared to CyaA, suggesting that the acylation of K860 and K983 stabilizes these three regions. These results are consistent with the HDX-MS analysis and indicate that acylations provide an apolar environment that favors the formation of the hydrophobic pockets in the AR domain, contributing to stabilizing and shaping CyaA.

**Figure 5:**
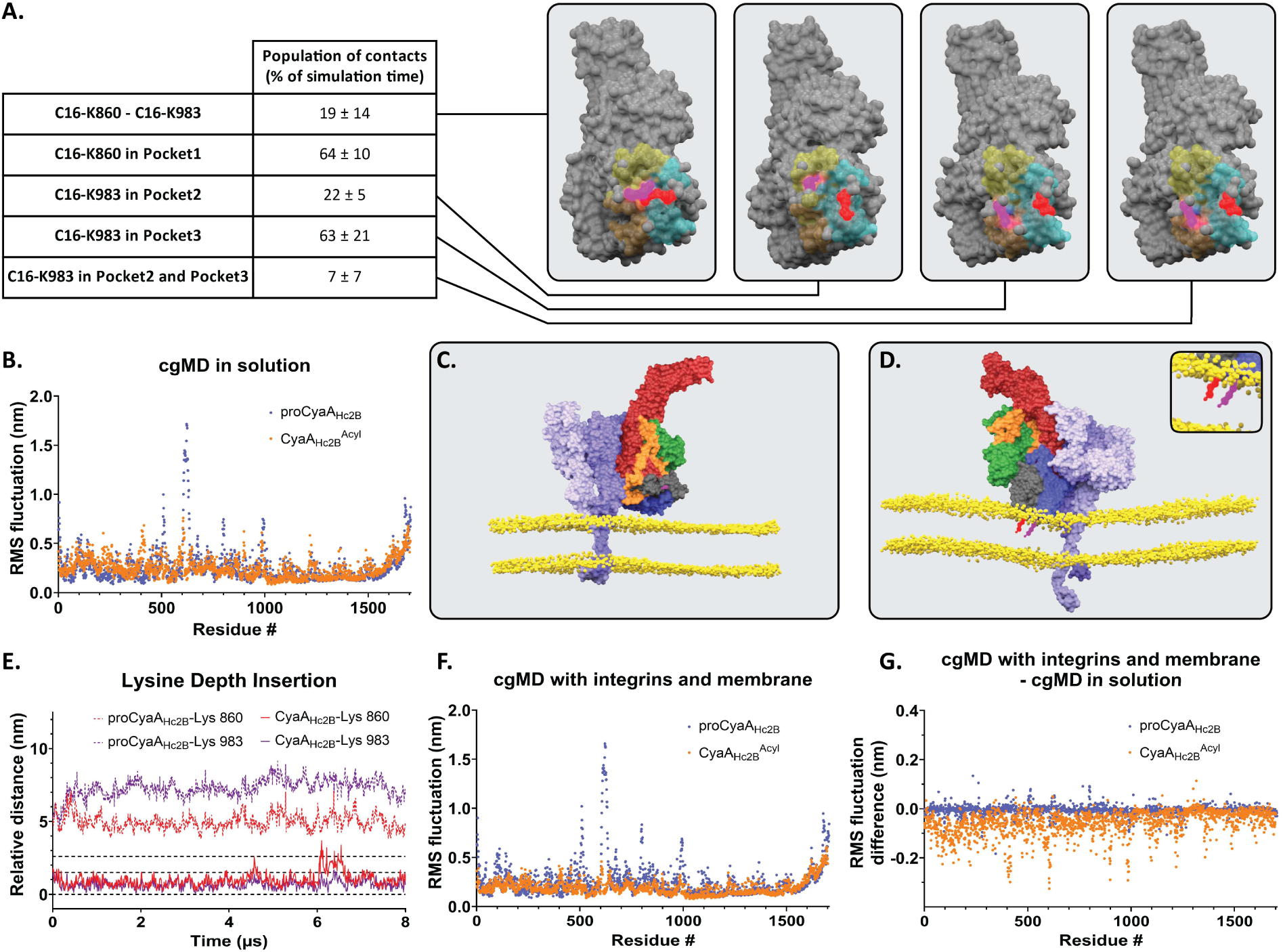
Molecular Dynamics simulations of proCyaA_Hc2B_ and CyaA_Hc2B_^Acyl^. A. Proportion of each location occupied by the two acyl chains during molecular dynamics of proCyaA_Hc2B_ and CyaA _Hc2B_^Acyl^ in solution: C16- K860 and C16-K983 interacting with each other, C16-K860 in pocket P1, C16-K983 in pocket P2, C16-K983 in pocket P3, and C16-K983 in pockets P2 and P3. Snapshots extracted from Supplementary Movie 3 with C16- K860 and C16-K983 in red and purple, respectively, and pockets P1, P2 and P3 in blue, yellow and orange, respectively. B. Root Mean Square Fluctuation (RMSF) of coarse-grained molecular dynamic (cgMD) simulations of the triplicate of proCyaA_Hc2B_ (blue) and CyaA_Hc2B_^Acyl^ (orange). C and D. Snapshot from MD simulations of proCyaA_Hc2B_ (C) and CyaA_Hc2B_^Acyl^ (D) in membrane containing CD11b/CD18 from Supplementary Movies 4 and 5. Both proteins are represented in sphere with ACD in green, TR in orange, HR in blue, AR in black and RD in dark red with lysine residues (proCyaA_Hc2B_) and acylated K860 and K983 (CyaA_Hc2B_^Acyl^) in red and purple spheres respectively. D, inset: zoom on the C16-K860 and C16-K983 in red and purple, respectively, of CyaA _Hc2B_^Acyl^ inserted in the lipid bilayer. Lipid headgroups are depicted in yellow; the acyl chains of the lipid bilayer and water molecules are omitted for clarity. Chimeric CD11b and CD18 integrins are represented in dark and light purple spheres, respectively. E. Depth insertion of lysine residues from proCyaA_Hc2B_ and CyaA_Hc2B_^Acyl^ into the lipid bilayer during the coarse-grained molecular dynamic simulations. The second bead of the lysine side chain was used as a reference to determine the depth of insertion in the membrane. The distances of the lysine residues are calculated from the center of mass of the lipid bilayer. During the cgMD simulations, C16-K860 (Cyan) and C16- K983 (blue) of CyaA_Hc2B_^Acyl^ are inserted into the membrane: on average, the K860 and K983 side chains are located at 1±0.5 nm and 0.7±0.3 nm from the center of the lipid bilayer, respectively. K860 (red) and K983 (green) of proCyaA_Hc2B_ remain far from the membrane, on average at 4.9±0.5 nm and 7±0.6 nm from the center of the lipid bilayer, respectively. The triplicate for each protein is shown in Supplementary Figure 3. Rolling means over 0.4µs are represented as lines over the data. See movie “Supplementary Movies 4 and 5”. Horizontal dashed lines at 0, 1.5 and 2.5 nm correspond to the center of the lipid bilayer (the interface between the *cis* and *trans* lipid leaflets), the interface between the acyl chain and the lipid headgroups, and the interface between the lipid headgroups and the solvent, respectively. F. RMSF of coarse-grained molecular dynamic (cgMD) simulations with integrins and membrane of the triplicate of proCyaA_Hc2B_ (blue) and CyaA_Hc2B_^Acyl^ (orange). G. RMSF subtraction between cgMD with integrins and membrane and cgMD in solution of proCyaA_Hc2B_ (blue) and CyaA_Hc2B_^Acyl^ (orange).

Finally, we performed coarse-grained MD simulations of proCyaA_Hc2B_ and CyaA_Hc2B_^acyl^ in the presence of a lipid bilayer containing the CyaA cell receptor, *i.e.*, the CD11b/CD18 integrin. Structural models of the complexes were generated by first aligning the AR/RD domains with the corresponding ones in the structure by Goldsmith *et al.*^28^ (PDB code: 7USL). Then, we assembled a chimeric integrin by coupling the ectodomain of CD11b and CD18^28^ to the transmembrane and intracellular segments of CD51/CD41-CD61^35,36^ (PDB codes: 2K9J and 1M8O) (**Extended Fig. 6**).

MD simulations of the complexes (**Fig. 5.C-G** and **Supplementary Movies 4** and **5**) revealed striking differences between the two systems. The acylation-induced elongated shape of CyaA appropriately positions the extremity of AR with acylated K860 and K983 close to the membrane surface. The two acyl chains in CyaA readily partition into the lipid bilayer, thus contributing to maintain the toxin in close proximity to the membrane (**Supplementary Movies 6** and **7**). In contrast, the AR domain in proCyaA is slightly more distant, and proCyaA moves away from the membrane surface (about 2 nm distance). Analysis of the insertion depth for the acyl chains confirms that both chains are responsible for firmly anchoring CyaA to the lipid bilayer (**Fig. 5.E and Supplementary** Fig.3). Additionally, RMSF analysis (**Fig. 5.F**) indicates that, consistent with behavior in solution, proCyaA remains more dynamic than CyaA. The presence of integrins and the membrane has a stabilizing effect on CyaA, whereas no such effect is observed for proCyaA, despite its persistent association with the integrins throughout the simulation (**Fig. 5.G**).

## Discussion

In this study, we report the first integrative structural models of the full-length adenylate cyclase CyaA toxin, and its non-acylated, inactive precursor, proCyaA. By combining complementary experimental techniques at different resolution with *in silico* integrative modeling, we circumvented long-standing challenges associated with the molecular mass, multi-domain architecture, structural flexibility, and aggregation propensity of CyaA. Our integrative approach sheds new light on the structure-function relationship of this multi-domain RTX toxin and reveals key mechanistic insights into the role of post-translational acylation in toxin folding and function.

A major outcome of this work is the generation of refined conformational ensembles for CyaA and proCyaA based on Cryo-EM single particle data combined with HDX-MS and SEC-SAXS experimental results. Most importantly, our findings highlight the importance of accounting for structural dynamics and conformational heterogeneity in multi-domain proteins. The failure of AlphaFold models to capture the structural diversity of CyaA underscores the utility of ensemble-based approaches such as our HDX-MS-guided coarse-grained MD simulations, reweighted by BioEM, *i.e.*, Hc2B pipeline developed in this study (**Extended Data** Fig. 1). This strategy allowed us to overcome limitations imposed by protein flexibility, preferential orientation on cryo-EM grids, and the lack of parameters for post translational acylation in current structure prediction algorithms. Beyond CyaA, our integrative pipeline offers a generalizable framework for the structural study of large and multi-domain proteins, characterized by structural dynamics and inter-domain flexibility, resistant to conventional high-resolution methods.

Our CyaA and proCyaA structural models show that acylation of K860 and K983 induces substantial conformational changes not only in the vicinity of the modified residues, but also at long distance in the ACD, HR, and AR domains while the C-terminal RD domain remains essentially unaffected (**Fig. 3 and 4**). More specifically, the post-translational acylation triggers the native folding and stabilizes conformations of HR and AR with hydrophobic pockets appropriately positioned and sized to accommodate the two acyl chains (**Fig. 4.D-E**). HDX-MS and coarse-grained MD simulations further show that the two acyl chains exhibit different behavior. The acylated K860 preferentially occupies the hydrophobic pocket P1, while the acylated K983 dynamically populates P2 and P3 (**Fig. 5.A** and **Supplementary Movie 3**). This dynamic exchange of the two acyl chains between the three pockets may facilitate the transfer of the acyl chains from the protein to the membrane upon CyaA binding to CD11b/CD18. In proCyaA, the absence of acylation leads to partially folded and relatively extended HR and AR (**Fig 4** and Extended Fig. 4 **and 5**). The stabilization of tertiary contacts between HR and AR in CyaA, and a more compact conformation of CyaA compared to proCyaA, strongly suggest that acyl chains act not only as membrane anchors, but also as critical trigger of toxin folding into its native and cytotoxic state.

Our MD simulations reveal that upon interacting with its cell receptor, CyaA adopts a conformation that optimally positions the acyl chains of K860 and K983 near the surface of the plasma membrane (**Fig.5.E** and **Supplementary Movies 4** and **6**). This conformation enables the coupling of receptor binding with membrane insertion of the acyl chains without requiring major structural rearrangements. This may account for the high affinity of CyaA for CD11b/CD18 expressing cells, and its quasi-irreversible association with myeloid cells^23^. This mechanism resembles a dual-lock system: protein-protein interactions between CyaA and the CD11b/CD18 receptor, together with the insertion of the two acyl chains into the plasma membrane, secure firm attachment of the toxin to target cells. Further membrane insertion of additional CyaA segments likely reinforces this interaction, contributing to the toxin’s quasi-irreversible, high-affinity binding to CD11b/CD18 expressing cells.

Taken together, our results support a model in which acylation serves as a conformational trigger that induces native folding and primes the toxin for membrane interaction, leading to the translocation of the catalytic domain across the plasma membrane. Future work should aim to capture membrane-inserted conformations of CyaA and describe the calmodulin-assisted ACD translocation across lipid bilayers^19^. Additionally, exploring whether similar acylation-dependent folding mechanisms exist in other RTX family toxins may uncover common structural principles of toxin folding, activation, and cell intoxication processes.

Overall, the results presented here have broad implications to understand the effect of post-translational acylation on proteins, which exhibit diverse biological functions, from bacterial toxins involved in host-pathogen interactions to eukaryotic proteins involved in the development of human diseases ^37,38^. It emerges that post-translational acylation contributes to protein folding in addition to its established role in membrane anchoring^39^. Such studies will expand our understanding of the functional significance of acylation in protein biology.

## Data Availability

SAXS data and atomic coordinates of AlphaFold models and CyaA_Hc2B_ and proCyaA_Hc2B_ are deposited in the Small Angle Scattering Biological Data Bank (SASBDB) with the following accession numbers: SASDXG6 and SASDXH6.

## Acknowledgments

This work was funded by the Agence Nationale de la Recherche (ANR 21-CE11-0014-01- 3DTransCyaA), Institut Pasteur (SPAIS/PTR 166-19, SPAIS/PTR 502-22, DARRI/Emergence) and CNRS. C.L. received funding from Institut Pasteur (SPAIS/PTR 166- 19) and ANR (ANR 21-CE11-0014-01-3DTransCyaA). S.E.H. was funded by a Pasteur-Roux-Cantarini fellowship from Institut Pasteur. G.S. was supported by the Institut Pasteur and Sorbonne University PhD program. A.A. was funded by a SPAIS/PTR 502-22 fellowship from Institut Pasteur. J.F. was funded by a Pasteur-Roux-Cantarini Tech fellowship and SPAIS/PTR 502-22 from Institut Pasteur. V.D. was funded by a Pasteur-Roux-Cantarini fellowship from Institut Pasteur. N.C. was supported by Institut Pasteur (DARRI/Emergence) and the ANR (ANR 21-CE11-0014-01-3DTransCyaA). L.M. acknowledges funding from the Institute National de la Santé et de la Recherche Médicale (INSERM). The HDX-MS platform was funded by the CACSICE Equipex (ANR-11-EQPX-0008). The NanoImaging Core was created with the help of a grant from the French Government’s “Investissements d’Avenir” program (EQUIPEX CACSICE “Centre d’analyse de systèmes complexes dans les environnements complexes”, ANR-11-EQPX-0008). Special thanks to SOLEIL for the provision of synchrotron radiation facilities, particularly SWING beamline at Synchrotron SOLEIL (St Aubin, France) for its support during X-ray diffraction data collection. The authors also thank the Molecular Biophysics and the Biological NMR and HDX-MS Technological Platforms and the Ultrastructural BioImaging, the Crystallography and the NanoImaging core Facilities at Institut Pasteur. The funders had no role in study design, data collection, analysis, decision to publish, or manuscript preparation.

## Methods

### Buffer

Buffer A: 20mM HEPES, 50mM NaCl, 2mM CaCl_2_, pH 7.4

Buffer B: 20mM HEPES, 6M Urea, 4mM CaCl_2_, pH 8.0

Buffer C: 20mM HEPES, 6M Urea, 50mM NaCl, pH 8.0

### Production and purification of proteins

Production and purification of proCyaA and CyaA were performed as previously described^40^. Production of CyaA and proCyaA monomers and their separation from multimers was performed on a BioSEC3 4.6 300 mm 300Å column (Agilent, 5190-2513). Briefly, a 200µL loading loop was washed by 500µL of buffer B then sequentially filled with 50µL of protein (CyaA or proCyaA in buffer C) then 100µL of buffer B. Elution was performed in buffer A. The procedure for the production of CyaA and proCyaA monomers is described elsewhere^24,31,40^.

### Static mass photometry

All mass photometry (MP) experiments were performed on a Refeyn TwoMP mass photometer at the PFBMI core facility (Institut Pasteur, Paris). Glass coverslips (24 x 50 mm, Menzel Gläser, VWR 630-2603) were washed with MilliQ water, isopropanol and MilliQ water. Silicone gaskets (Grace Bio-Labs, GBL103280) were rinsed in MilliQ water and isopropanol and dried under a nitrogen stream. Gaskets were installed on the coverslips. Wells were filled with a drop of buffer (18 μL of 20mM HEPES, 150mM NaCl, 2mM CaCl_2_, pH 7.4) first, then 2 μL of protein samples were diluted to 10 nM in the buffer drops. MP data were acquired and processed with AcquireMP and DiscoverMP, respectively.

### Analysis of cAMP production by CyaA and proCyaA in reporter cell line

Stable monoclonal NCI-H292 cells expressing a luciferase-based biosensor for real-time cAMP detection, called cAMP-GloSensor cells^32^, were seeded at 2.0 x 10^4^ cells per well in a 96-well plate (Corning, reference 3610). Cells were incubated for two days at 37°C and 5% CO_2_ in RPMI medium supplemented with 10% fetal bovine serum (FBS). Prior to intoxication, cell medium was removed, and cells were incubated in the dark at room temperature for one hour in phenol red-free RPMI medium containing 5% GloSensor™ cAMP Reagent (Promega, E1290), 2 mM CaCl_2_ and 0.5% BSA. The cells were then intoxicated with different concentrations of CyaA or proCyaA monomers, ranging from 1.24 to 0.037 nM and luminescence emission was monitored at 24°C using an automated plate reader (TECAN Spark^®^ 10M) and an integration time of 1s.

### SEC-SAXS

Size-exclusion chromatography (SEC) coupled to small-angle X-ray scattering (SEC-SAXS)^41^ measurements were carried out on the SWING beamline of the SOLEIL Synchrotron Radiation Facility (Saint-Aubin, France). **Supplementary Table 3** gives all the experimental details in accordance with BioSAXS publication guidelines^42^. For proCyaA and CyaA, measurements were performed using a size-exclusion HPLC BioSec3 4.6*300 300Å column (Agilent, 5190-2513) online with the SAXS measuring cell, a 1.5 mm diameter quartz capillary contained in an evacuated vessel^43^. Samples were prepared in buffer A. Briefly, monomers of proCyaA and CyaA were produced and then concentrated up to 17µM. Afterwards, sample of 50 μL were loaded onto the SEC column. Scattering of the elution buffer before void volume was recorded and used as buffer scattering for subtraction from the protein patterns. Successive frames of 1s were recorded. The elution flow of 0.300 mL/min ensured that no protein was irradiated for more than 0.4s. Primary data reduction was performed using FOXTROT, the SWING in-house software. This yielded azimuthally averaged scattering intensities I(q) put on an absolute scale (cm^-1^ units) using water scattering, where q is the momentum transfer (q = 4π sinθ/λ, where 2θ is the scattering angle and λ the wavelength of the X-rays). Data were subsequently processed using the program package PRIMUS^44^ and RAW^45^. The forward scattering I(0) and the radius of gyration (Rg) were evaluated using the Guinier approximation^44^. Frames over the elution peak were individually analysed before averaging the appropriate subset that yielded identical I(q)/c profiles with US-SOMO^46,47^. For proCyaA, due to its high aggregation propensity, two scattering profiles measured at different initial concentrations (5.3 and 16.1 µM) were merged. The scattering profile of the low proCyaA concentration contains a reduced contribution of aggregates at small q but is noisier at higher q than the scattering profile measured with proCyaA at 16.1 µM (**Extended Data** Fig. 3**.B**). The distance distribution function P(r) was determined using the indirect Fourier transform method as implemented in the GNOM program^48^.

### Negative staining transmission electron microscopy

Four µL of CyaA monomers at 20 nM was applied to carbon-coated copper grids, negatively stained with 2% uranyl acetate, and observed with a transmission electron microscope Tecnai T12 (FEI). The images were digitally recorded, using an ultrascan camera (Gatan) with the TIA software. (**Supplementary** Fig. 1)

### Cryogenic electron microscopy

Cryo-EM samples were prepared in 20 mM HEPES, 50m M NaCl, 2mM CaCl_2_, pH 7.4. They were applied to glow-discharged C-flat carbon grids 1.2-1.3 mesh 300 and vitrified using a Vitrobot Mark IV (Thermo Fisher). Following this treatment, the grids were stored in liquid nitrogen until analysis. Grids were then imaged on a Glacios (200 keV, Thermo Fisher) in electron counting mode with a raw pixel size of 0.95 Å. For CyaA and proCyaA, respectively 21,682 and 5,258 raw movies were collected using the EPU software with a defocus ranging from -1 to -4µm with ∼ 40-50e per Å² at ∼10e/px/s. Raw movie frames were treated using Cryosparc software. They were aligned and dose-weighted using “Patch Motion Correction”. CTF parameters were estimated using “Patch CTF Estimation”. Movies were then filtered and final ensembles of 8,366 and 3,237 treated movies were obtained. The raw dimension of the proteins for single-particle picking was defined based on SEC-SAXS results, *i.e.*, an elliptical blob of 50 by 180Å and 50 by 210Å for CyaA and proCyaA, respectively. Based on these elliptical blob dimensions, 3,348,038 and 295,415 particles were picked for CyaA and proCyaA, respectively. They were then subjected to several rounds of reference free 2D classifications, removing bad classes between each round which led to a total of 647,213 and 110,173 single particles of CyaA and proCyaA, respectively. Several *Ab initio* models were obtained and subsequent optimization and refinement produced 5.5-8 Å resolution density maps for both proteins^49^.

### Alphafold modeling

Five models of proCyaA have been generated with AlphaFold2.2^50^. They show four well-folded domains (AC, HR, AR and RD) and one unfolded domain (TR). The models are quite extended (R_g_=53-57 Å, D_max_=193-211 Å). AlphaFold model 3 (AFSM#3) is in best agreement with SEC-SAXS data and with previously published AUC and SEC-TDA data (**Supplementary Table 1 and 2**) and has been therefore used for further analysis. The average per-residue pLDDT and associated standard deviations of this model were 76.9/17.2, 24.7/3.5, 35.7/21, 88.8/7.7 and 88.4/13.8 for the AC, TR, HR, AR and RD domains, respectively. Two independent runs were performed with AlphaFold3 for both forms of the toxin (4 runs in total), including 36 calcium ions (for proCyaA and CyaA) and acylated lysine residues as PTMs (only for CyaA). Hydrodynamic parameters showed discrepancies between each run with mean R_g_ of 49 and 61 Å for CyaA, 47 and 59 Å for proCyaA and mean D_max_ of 180 and 222 Å for CyaA, 186 and 224 Å for proCyaA. No improvement was therefore obtained with AlphaFold3 compared to AlphaFold2.

### All-atom molecular dynamics simulations of proCyaA and CyaA

Calcium ions (Ca^2+^) required for the stability of the RTX blocks of the receptor-binding domain (RD) were added to AFSM#3 using a set of X-ray structures of individual CyaA RTX blocks: the pdb 7RAH for RD blocks I-III^51^, the pdb 6SUS for RD block IV^52^ and the pdb 5CVW for RD block V^52^. After superposition of an X-ray structure onto the corresponding RD block of AFSM#3, the atomic coordinates of the Ca^2+^ ions were transferred to the AF model only when a carboxyl group was found in sufficient proximity for coordination, for a total of 36 ions. To ensure stability of the Ca^2+^-coordinated regions during the MD simulation, harmonic distance restraints centered at a canonical distance of 2.4 Å^53^ were added between the Ca^2+^ ions and coordinating oxygens.

ChemDraw was used to design the acylated lysine containing an acyl chain (a palmitoleic acid, C16:1Δ9, *i.e.*, (9*Z*)-hexadec-9-enoic acid, CAS registry number: 373-49-9) composed of 1 COO, 7 CH_2_, 1 C=C with the H in trans between positions 9 and 10, 5 CH_2_, and a terminal CH_3_. Pymol (The PyMOL Molecular Graphics System, Version 3.0 Schrödinger, LLC.) was used to superimpose an acylated lysine to residues K860 and K983 in the AFSM#3, calcium-loaded model of proCyaA. Each acyl chain was connected to the lysine residue by replacing an HZ atom of K860 and K983. A new residue, ACK, was added to the CHARMM36 forcefield^54^ importing bonded and non-bonded force field parameters from the CHARMM36 lipid. CHARMM-GUI v. 3.7^55–57^ was used to prepare both proCyaA and CyaA topologies.

GROMACS v. 2019.4^58^ was used to set up a system containing the calcium-loaded proCyaA and CyaA models described above. The proteins were first embedded in a triclinic box with dimensions such as the closest protein atom were 12 Å away and then immersed in water molecules. The total charge was neutralized at a concentration of 0.15M by adding 336/305 and 303/340 chloride/potassium ions for proCyaA and CyaA, respectively. CHARMM36m and mTIP3P^59^ force fields were used for protein and solvent molecules, respectively. The Particle Mesh Ewald approach^60,61^ was used for long-range electrostatic with a cutoff of 12 Å. A time step of 2 fs was used together with LINCS constraints on H-bonds^62^. Each system was first subject to 100,000 steps of steepest descent energy minimization, followed by 1 ns of equilibration in the NVT ensemble using a Nose-Hoover thermostat to maintain the temperature at 310.15 K. Production simulations were performed in the NPT ensemble using the Parrinello- Rahman barostat^63^ for 1.5 µs and 1.1 µs for proCyaA and CyaA, respectively. Models were clustered using the *gromos* algorithm implemented in GROMACS, using the Cα-RMSD as metrics with cutoff equal to 0.3 nm.

### Coarse grained molecular dynamics simulations of CyaA and proCyaA

We used coarse-grained (CG) molecular dynamics simulations to sample the conformational landscape of CyaA and ProCyaA. The atomic-resolution AFSM#3 model was first converted to a Martini 3^64^ CG representation^65,66^ using martinize2^67^. In standard Martini simulations, the secondary and tertiary structures of proteins are restrained using an elastic network, limiting exhaustive sampling of the conformational landscape. We therefore selectively removed these restraints using experimental information on the stability of protein secondary structure obtained by HDX-MS experiments. Residues with higher deuterium exchange rate were considered to be more dynamic, and the corresponding elastic restraint network was removed. For each protein, the CG model was then placed in a cubic simulation box with an edge length of 23nm and immersed in an aqueous solution with 0.15M NaCl.

All simulations were performed with GROMACS v. 2020.5^58^ and the Martini 3 forcefield^64^ with standard parameters. For long-range electrostatics interactions, we used the reaction-field method with a screening constant of 15 and a cutoff of 1.1 nm. Van der Waals interactions were switched off at 0.9 nm and cutoff at 1.1 nm. Each system was first energy-minimized using the steepest gradient descent algorithm, followed by 10 ns of NVT equilibration using the Bussi-Donadio-Parrinello thermostat at 300 K with a time step of 1 fs and position restraints on the protein. Next, we performed 10 ns of NPT equilibration with no position restraints, using a 1 fs timestep with the Berendsen barostat^68^ and the Bussi-Donadio-Parrinello thermostat^69^ at 300 K. Production simulations were performed in the NPT ensemble with a 20 fs timestep for 10 µs using the same barostat and thermostat as in the NPT equilibration. Structural clustering of the resulting 1,000 conformations was carried out using the GROMACS cluster linkage algorithm with an RMSD cutoff of 8 Å, resulting in 127 and 121 distinct protein conformations for proCyaA and CyaA, respectively.

### Ensemble refinement using cryo-EM single-particle images

To refine the conformational ensembles generated by the HDX-MS Martini simulations described above, we employed BioEM^70,71^. Image-model likelihoods were calculated for each protein model-single particle pair for the 121 models and 3,255,453 images of CyaA and the 127 models and 162,200 images for proCyaA. The models input to BioEM were generated by converting the backbone atoms of the clustered coarse-grained models of CyaA and proCyaA to Cα carbon atoms. This representation offers sufficient accuracy, as BioEM describes protein density as a series of spheres centered on the Cα carbon atoms. Once the likelihoods 𝑃*_iω_*” were calculated for each model-particle pair, we estimated the population of each model 𝑤_i_ by maximizing the BioEM posterior probability^71^ of observing models 𝑀 given the set of single-particle images 𝛺:

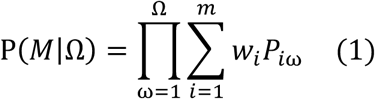

To identify the set of (normalized) weights 𝑤_i_ that maximize P(𝑀|Ω), we first defined the associated BioEM energy function as:

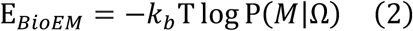

where 𝑘_b_T was set to 1.0 and then sampled the weights using a Monte Carlo (MC) approach. To ensure efficient sampling, we used metadynamics^72^ to add Gaussian repulsive potentials on the energy collective variable^73^. Gaussians of height equal to 1000 and width equal to 100 were added every 1 MC steps. For each system, we performed 5 independent MC simulations and stored the set of weights that resulted in the minimum value of E_BioEM_ (i.e. maximum posterior probability) in each run. Finally, we computed our best estimate of the weights as the average of the weights that minimized E_BioEM_in each replicate MC simulation.

### Calculation of theoretical SAXS profiles and comparison with experimental data

We utilized Pepsi-SAXS^74^ to compute theoretical SAXS profiles from individual models as well as structural ensembles. To do this, the coarse-grained models of CyaA and proCyaA were first converted to atomic-resolution models using the backward.py script provided by Wassenaar *et al.* The resulting models were then optimized using SCWRL4^75^ and used for SAXS calculations using Pepsi-SAXS (**Fig. 3**). The calculation of theoretical SAXS curves for structural ensembles followed the procedure defined in Pesce *et al.*^76^, which is briefly summarized here. We first computed, for each ensemble conformer, a theoretical profile 𝐼_th_(𝑞) using a fixed scale factor of 1 and no offset. For each conformer, we scanned 31 values of the hydration shell contrast *δρ* between 3.34*10^-5^ and 3.34*10^-2^ eÅ^-3^, and 11 values of the effective atomic radius r_0_ between 1.45 and 1.77 Å. For each choice of parameters, an ensemble-averaged profile 𝐼_th_(𝑞) was then computed across all conformers. Model weights for the calculation of the weighted average in the standard and HDX-MS Martini ensembles were determined as the normalized cluster size; for the BioEM ensemble, the weights determined by maximizing the posterior in Eq. 1 were used. An optimal scale factor *f* and offset *c* were then computed for each *δρ* and r_0_ pair to minimize the reduced χ^2^ with respect to the experimental SAXS profile 𝐼_exp_(𝑞):

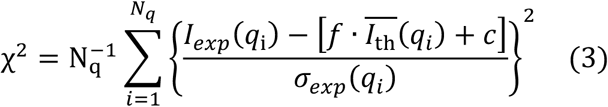

where 𝑁_q_is the number of data points in the experimental SAXS curve and 𝜎_exp_(𝑞) the experimental error associated to each data point. Finally, the *δρ* and r_0_ pair that resulted in the minimum value of χ^2^across all the parameters tested was selected. To compute theoretical SAXS profile from individual models, the same procedure was used, except for the ensemble-average calculations.

### Pocket identification

To identify potential pockets that could accommodate the acyl chains connected to K860 and K983, we employed *mkgridXf* ^77^. The default settings for mkgridXf were employed using the most populated models after sidechain optimization using SCWRL4. To determine the pocket hydrophobicity, we labelled each natural amino acid as hydrophobic (I,L,F,M,C,Y,W), hydrophilic (P,H,E,Q,N,R,K,D), or neutral (T,S,G,A,V). We then computed the number of hydrophobic residues within the pocket as identified by *mkgrifXf* and normalized by the total number of pocket residues.

### ProCyaA and CyaA integrin complexes generation

We first made a structural alignment based on integrin interaction with RD to replace N-RTX and AR from 7USL by our models (CyaA_Hc2B_^acyl^ or proCyaA_Hc2B_) and the ScFv in the initial pdb was deleted. Based on https://pdb101.rcsb.org/motm/134, inactive conformation of integrin CD51/CD41 - CD61 was reconstructed (pdb codes 1JV2, 2K9J and 1M8O). Through structure and sequence alignments, extracellular parts of this reconstructed integrin were substituted by the extracellular parts of integrins CD11b - CD18 present in 7USL pdb structure. This gave a chimeric integrins with extracellular CD11b-CD18 and a transmembrane-intracellular CD51/CD41-CD61. Then, a second structural alignment was made to obtain CyaA_Hc2B_^acyl^ and proCyaA_Hc2B_ in complex with the chimeric integrin. Finally, a lipid bilayer was added based on the transmembrane part of the chimeric integrins.

### Molecular dynamics simulations in solution and in lipid membranes: system setup and simulation parameters

Acylations (C16:1Δ9, *i.e.*, (9*Z*)-hexadec-9-enoic acid, CAS registry number: 373-49-9) on K860 and K983 were added using pymol on CyaA_Hc2B_ which gave CyaA_Hc2B_^acyl^. We then used coarse-grained (CG) molecular dynamics simulations to characterize the interaction of CyaA and ProCyaA with lipid membranes and integrin receptors. All the simulations were carried out using the GROMACS software (version 2022)^58^. Atomistic structures of proCyaA, CyaA, and chimeric integrin receptors^78^ were transformed into Martini CG models with the martinize2^67^. Coarse-grained (CG) molecular dynamics simulations were performed with the Martini^65,64^ force field (v3.0), which still uses the same definition of protein backbone as in the original Martini protein force field^66^. The secondary and tertiary structure of all proteins were fixed using an elastic network of harmonic bonds. For CyaA, parametrization of two palmitoleic acyl chains attached to lysines (K860 and K983) was based on palmitoyl chains in lipids, and the link between the lysines and the palmitoleic chains was based on standard Martini procedures (see https://vermouth-martinize.readthedocs.io/en/latest/tutorials/6_adding_residues_links/index.html). For the CyaA:Integrins complex (built as described above), the elastic networks covered each individual protein but not the complex, i.e., no restraints were imposed on the relative positions of the proteins nor on the binding between CyaA and integrins.

The complex was inserted in a large (40 x 40 nm) membrane using the insane.py script^79^, with the following composition: POPC:DPSM:CHOL:POPE 35:30:30:5 (with concentrations expressed as molar ratio) for the outer leaflet and POPC:POPE:CHOL:POPS:DPSM 23:27:28:17:5 for the inner leaflet. The protein:membrane system was solvated using standard Martini water and ions, with a 150 mM concentration of Na^+^ and Cl^-^ ions. Non-bonded interactions were calculated using standard Martini parameters, using the Verlet cut-off scheme, as detailed above for the CG simulations of CyaA in water. The solvated system was first energy minimized for 1000 steps, and then equilibrated with position restraints on the protein (force constant of 4,000 kJ.mol^-1^.nm^-2^ on all protein backbone beads), in the NPT ensemble using the Bussi-Donadio-Parrinello ^80^ thermostat (temperature of 310 K, tau_T_=1 ps) and c-rescaling barostat^81^ (pressure of 1 bar, tau_P_=12 ps) for 100 ns. The integration time step was 10 fs, to increase numerical stability. For each system we carried out a set of two simulation replicas (10 µs each) to assess the stability of the protein complex. Similar procedure was applied to prepare the system for proCyaA and CyaA monomers in water and ions (except for insane.py that was not required) in a 30 x 30 x 30 nm box with the same concentration of ions. A time step of 20fs was used for the production run. For each and every configuration a set of two 5µs replicas was produced to extract information on the conformation of the protein and side chains using previous equilibration and production parameters.

### Simulation analysis

A first qualitative visual inspection was first done using VMD^82^. The root-mean-square fluctuation (RMSF) of atomic positions were calculated using GROMACS^58^ tool gmx rmsf. To assess the role of lysine palmitoylation in the insertion mechanism, quantitative analysis of the trajectories was performed using MDAnalysis^83^ to measure the insertion of the acyl chains (in CyaA) and lysine side chains (in proCyaA) in the membrane; the second bead of the lysine side chain was used as a reference to assess the insertion of the residues.

---

**Extended Data Fig. 1.**
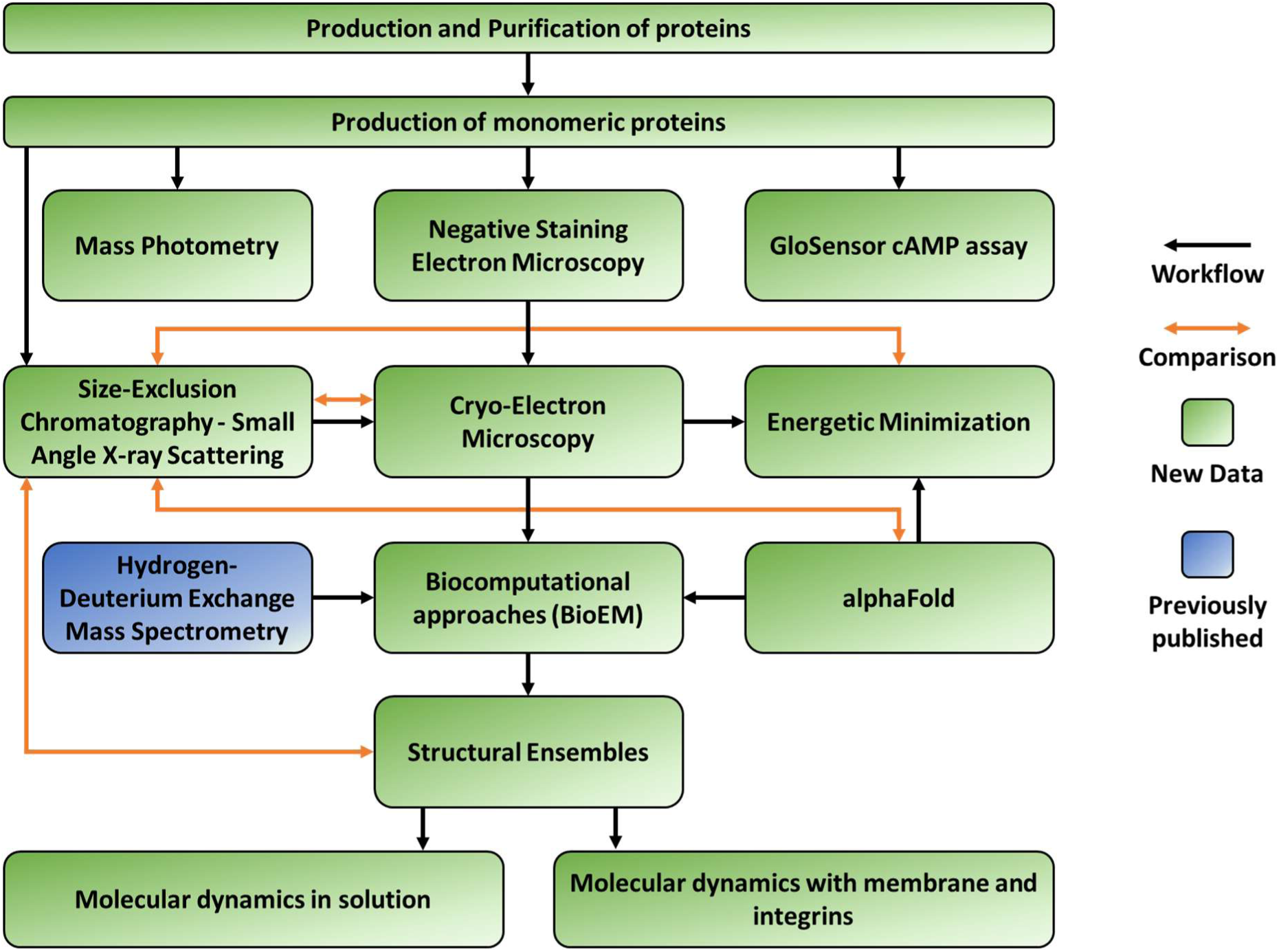
Pipeline of the integrative structure determination of proCyaA and CyaA. Schematic representation of the experimental and computational pipeline used to determine individual structures and structural ensembles of proCyaA and CyaA. Blue boxes represent data from previous works; green boxes indicate newly generated data. Black arrows denote the experimental workflow, and orange double-arrows indicate comparisons of structural models with in-solution SEC-SAXS profiles. Following protein production and purification, monomeric species were analyzed using mass photometry, negative staining electron microscopy, and GloSensor cAMP assays to ensure the presence of monomeric and functional proteins. In solution low-resolution structural insights were obtained through size-exclusion chromatography coupled to small-angle X-ray scattering (SEC-SAXS). Single- particle cryo-electron microscopy (cryo-EM) and hydrogen-deuterium exchange mass spectrometry (HDX-MS) were then integrated with computational methods including AlphaFold predictions, atomistic and coarse-grained molecular dynamics, and refinement of structural ensembles with single- particle images. The resulting models of CyaA and proCyaA were validated by SAXS and used to perform coarse-grained molecular dynamics simulations both in solution and in the context of a lipid bilayer and protein partners.

**Extended Data Fig. 2.**
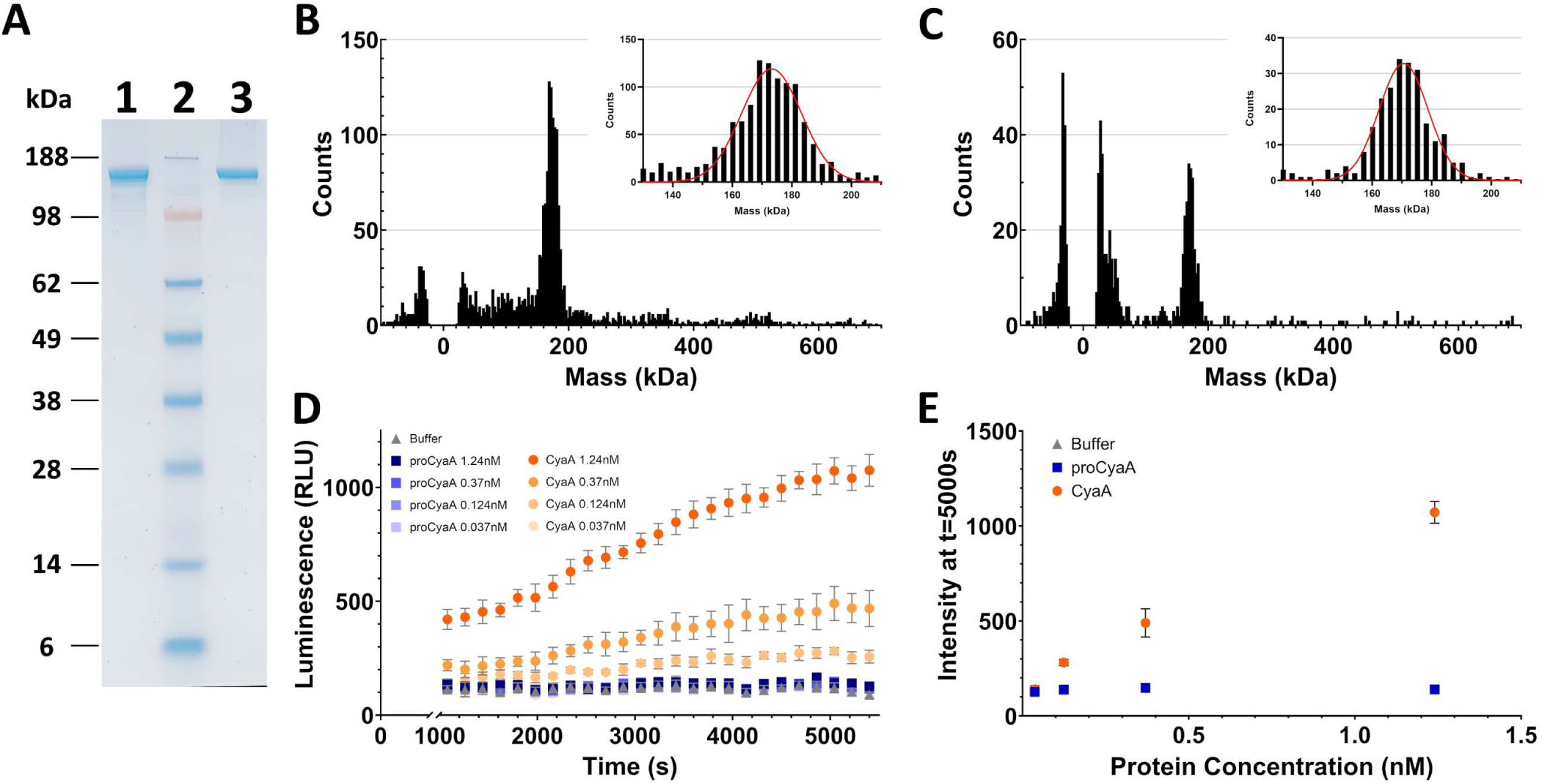
Stochiometric, structural and functional characterization of proCyaA and CyaA. **A.** SDS-PAGE analysis of proCyaA (lane 1) and CyaA (lane 3) at the end of the purification procedure, before refolding the proteins by molecular confinement. Molecular mass standards are reported in lane 2. B-C. Mass photometry analyses of monomeric proCyaA (B) and CyaA (C) samples performed at 10 nM in 20 mM HEPES, 150 mM NaCl, 2 mM CaCl_2_, pH 7.4. Insets focus on the 130– 210 kDa range. The positive and negative peaks at circa ±40 kDa likely correspond to the dissociation from the coverslip (peak at -40 kDa) and the re-adsorption on the coverslip (peak at +40 kDa) of the CyaA catalytic domain (ACD). D. Luminescence emission of cAMP-GloSensor cells following intoxication with CyaA. Luminescent reporter cells were intoxicated with different concentrations of CyaA monomers (orange), proCyaA monomers (blue) or buffer (grey line) ranging from 1.24 (darkest color) to 0.037 nM (lightest color). Luminescence intensity represents intracellular cAMP levels. E. Luminescence intensity of cAMP-GloSensor cells after 5000 seconds of intoxication with the various protein concentrations.

**Extended Data Fig. 3.**
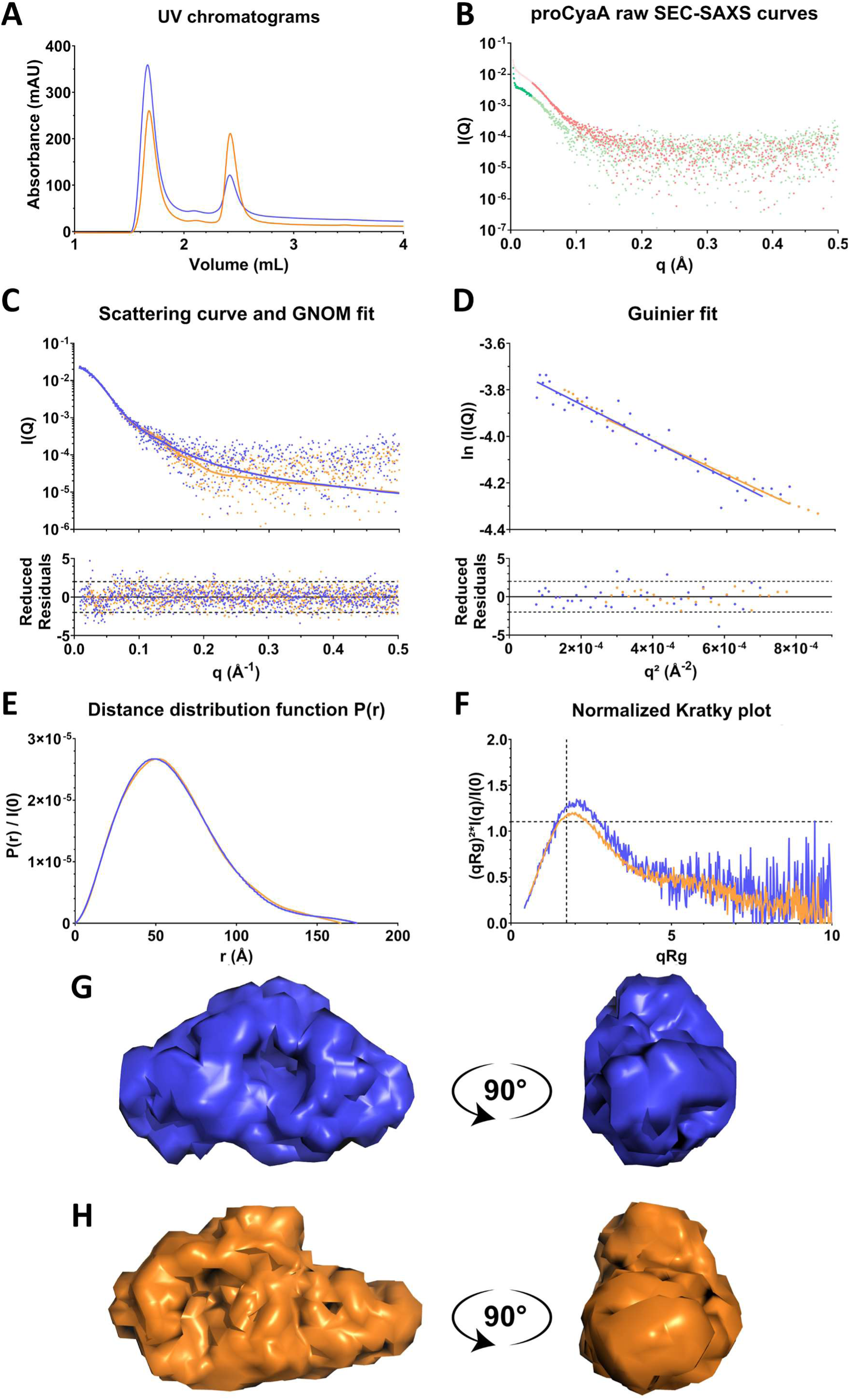
Size Exclusion Chromatography Coupled with Small Angle X-Ray Scattering (SEC-SAXS) of proCyaA and CyaA. A. UV profiles of proCyaA (blue) and CyaA (Orange) samples analyzed on a BioSec3 4.6*300 300Å column in 20mM HEPES, 50mM NaCl, 2 mM CaCl_2_, pH 7.4. Monomeric proteins were concentrated up to ∼16µM (∼3g/L) prior to SEC-SAXS analysis. This concentration led to the formation of multimeric species as shown in the elution profiles. The first peak (eluting at ∼1.7mL, in the dead volume) corresponds to multimers and the second peak (eluting at ∼2.4mL) to monomers. B. proCyaA raw SEC-SAXS curves obtained at 0.95 mg/mL (Green) and 2.86 mg/mL (Red). The final proCyaA curve is composed of SAXS data from the low (dark green) and high (dark red) concentration samples. C. Top: Experimental SAXS intensities for CyaA and proCyaA (dots), and GNOM fit (lines). Bottom: Distribution of reduced residuals corresponding to each fit. D. Top: Guinier fit over small q-range. Bottom: Reduced residuals associated with each Guinier fit. E. P(r) distance distribution of each form of the toxin. F. Normalized Kratky plot. The dashed vertical line at √3 and horizontal line at 1.104 indicate the characteristic peak for globular proteins. G-H. DENSS models of (G) proCyaA and (H) CyaA. The proCyaA and CyaA data are available on SASBDB with the following accession code: SASDXH6 and SASDXG6, respectively.

**Extended Data Fig. 4.**
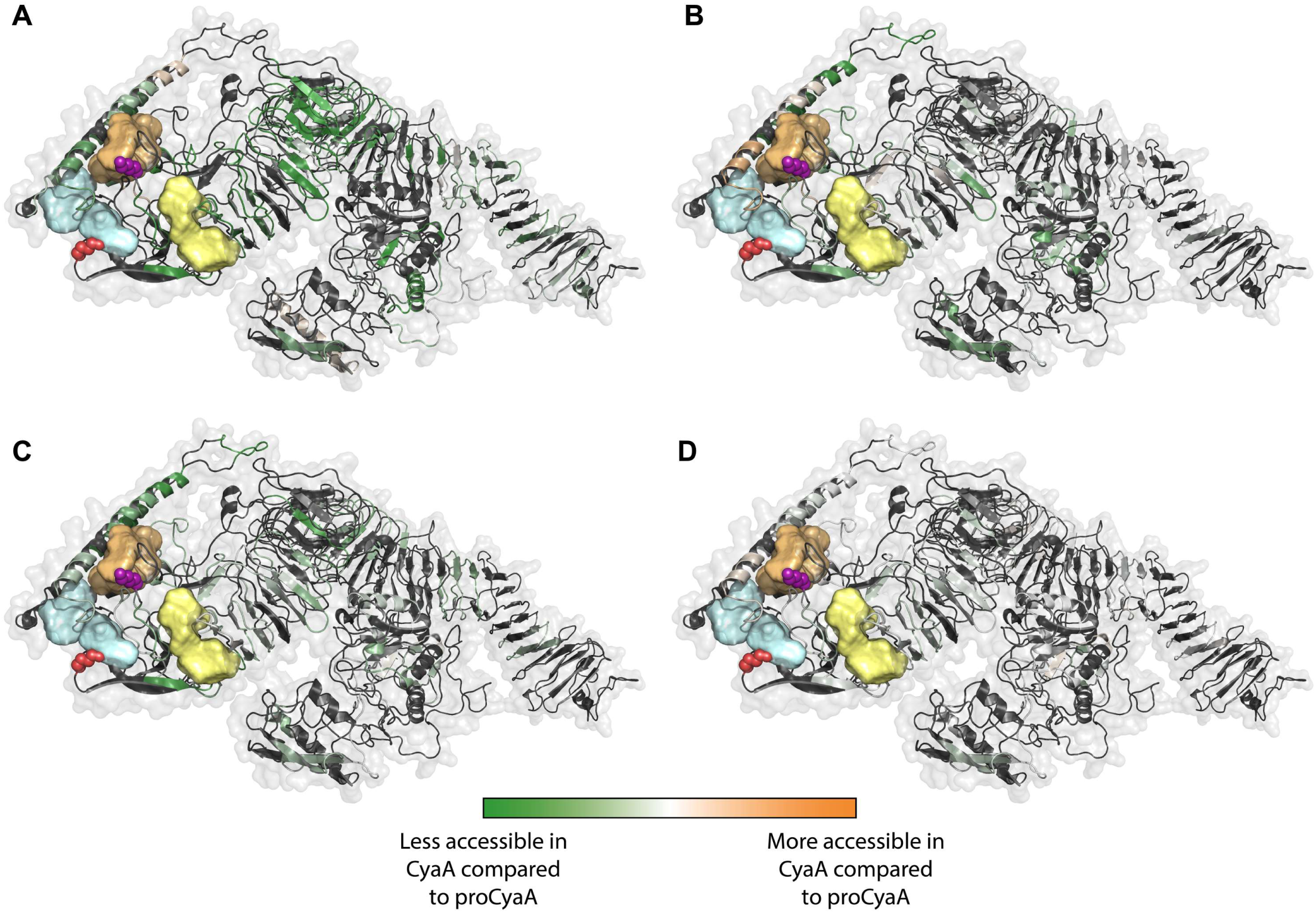
Effect of acylation on CyaA folding at different urea concentrations followed by HDX-MS. HDX-MS uptake differences between CyaA and proCyaA at different concentrations of urea (A. 0M, B. 1.6M, C. 3M, D. 6M urea) mapped on CyaA_Hc2B_ using a cartoon representation with lysine residues K860 and K983 visualized as red and purple spheres, respectively. Pockets are represented in blue (Pocket 1, P1), yellow (P2) and orange (P3). The color scale ranges from green (less accessible in CyaA compared to proCyaA) to white then orange (more accessible in CyaA compared to proCyaA). Protein regions not covered by HDX-MS are shown in black.

**Extended Data Fig. 5.**
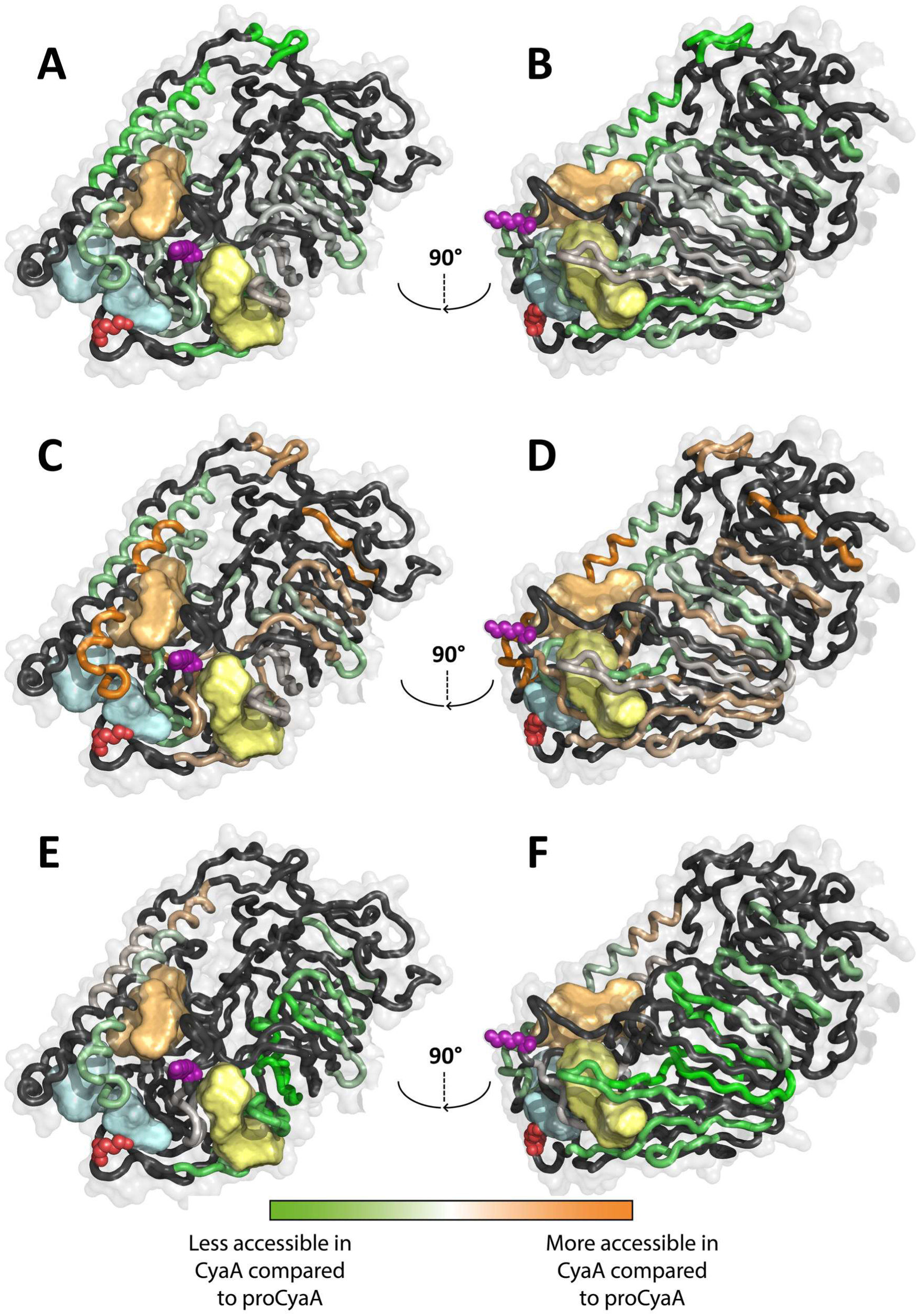
Impact of acylation on proCyaA and CyaA folding followed by HDX-MS as a function of urea concentration. The HDX-MS uptake differences are mapped from residue 496 to 1141 of CyaA_Hc2B_. Models are represented in ribbon with lysine residues K860 and K983 visualized as red and purple spheres, respectively. Pockets are represented in blue (Pocket 1, P1), yellow (P2) and orange (P3). HDX-MS uptake differences from 6M to 3M urea (A. and B.), from 3 to 1.6M of urea (C. and D.) and from 1.6 to 0M of urea (E. and F.). Scale color ranges from green (less accessible in CyaA than in proCyaA, while urea concentration decreased) to light grey then orange (more accessible in CyaA than in proCyaA, while urea concentration decreased). Black regions indicate residues not covered by HDX-MS. The HDX-MS uptake differences between CyaA and proCyaA were compared from 6 to 3 M urea, from 3M to 1.6M urea and finally from 1.6M urea to the urea-free buffer. The data show that from 6 to 3M urea, the location of the acyl chains in the blue (K860) and orange (K983) pockets notably favor the refolding of the HR helical hairpin of CyaA compared to proCyaA (highlighted in green in A. and B.). The structural elements underneath the yellow pocket in A. and B. are moderately stabilized by the presence of the acylations in CyaA compared to proCyaA. From 3 to 1.6M urea, parts of HR and AR of proCyaA are stabilized (segments colored in orange in C. and D.). In particular, the long helices of HR are stabilized in proCyaA from 3 to 1.6 M urea, while this stabilization effect was observed from 6 to 3M urea in CyaA (A. and B.). This suggests that this HR region can refold in both CyaA and proCyaA, however, the refolding occurs at higher urea concentration in CyaA, indicating that the acylation triggers the refolding of HR and induces a stronger stabilization of this region, compared to proCyaA (in which the refolding and stabilization occur but only between 3 and 1.6M urea). The most important stabilization effect is measured from 1.6 M to 0M urea for CyaA in the acylation region (green traces in E. and F. around the yellow pocket). This suggests that the K983 acylation located in the yellow pocket is triggering the folding and stabilization of AR. This result may also explain why AR is only partially folded in proCyaA (Fig. 3.B).

**Extended Data Fig. 6.**
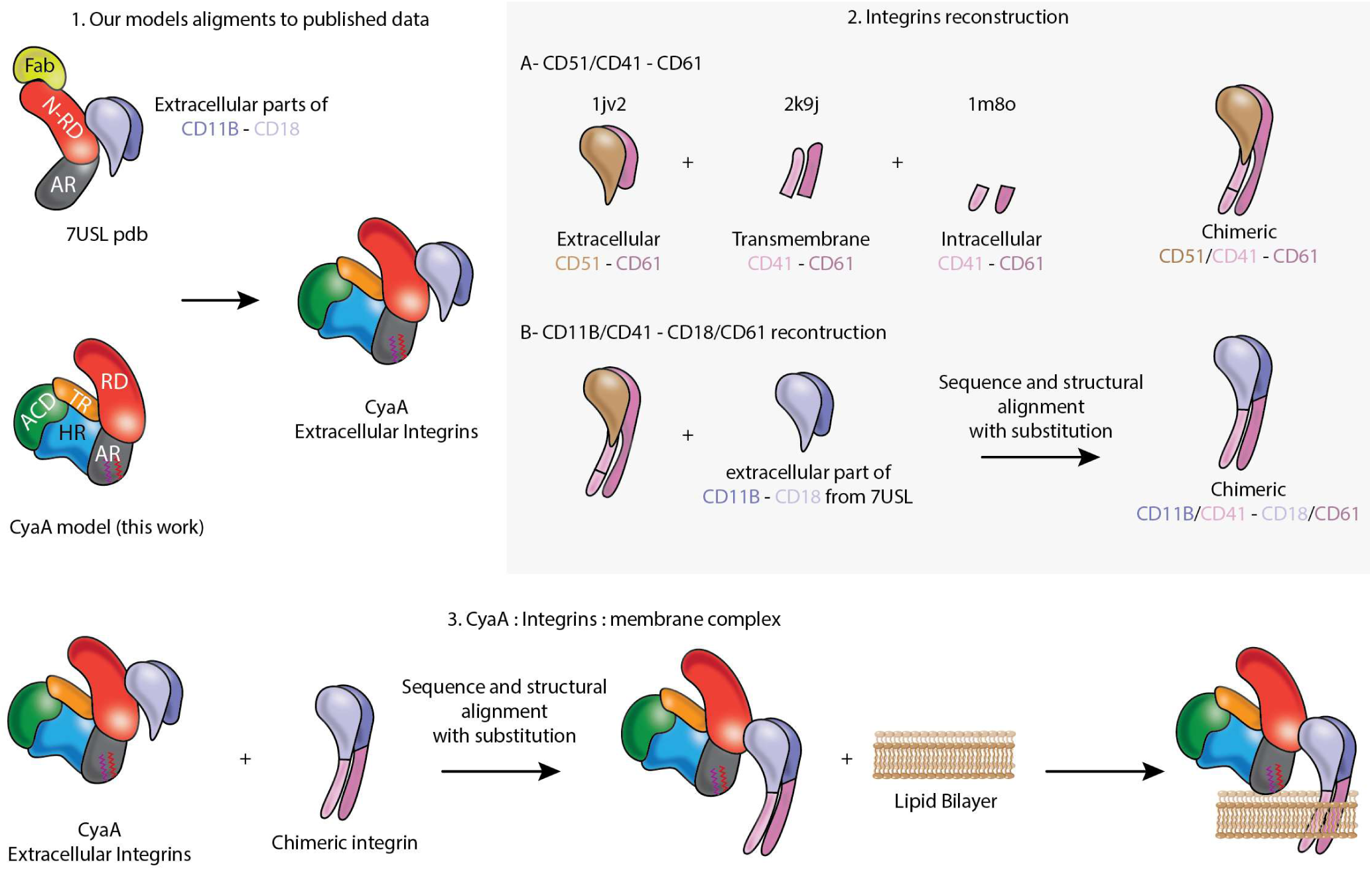
Assembling the model of the CD11b-CD18 CyaA complex. We first made a structural alignment based on integrin interaction with RD to replace N-RTX and AR in PDB 7USL by our model (CyaA_Hc2B_^Acyl^ or proCyaA_Hc2B_) and the ScFv in the deposited structure was deleted. Based on https://pdb101.rcsb.org/motm/134, an inactive conformation of integrin CD51/CD41 - CD61 was reconstructed (PDB codes 1JV2, 2K9J, and 1M8O). Through structure and sequence alignments, extracellular parts of this reconstructed integrin were substituted by the extracellular parts of integrins CD11B - CD18 present in PDB 7USL. This procedure resulted in a chimeric integrin with extracellular CD11B-CD18 and a transmembrane-intracellular CD51/CD41-CD61. A second structural alignment was then made to obtain CyaA_Hc2B_^Acyl^ and proCyaA_Hc2B_ in complex with the chimeric integrin. Finally, the transmembrane region of the chimeric integrins was used to adjust the position of the lipid bilayer.

## Bibliography

1. Carbonetti, N. H. Pertussis Toxin and Adenylate Cyclase Toxin: Key Virulence Factors of *Bordetella Pertussis* and Cell Biology Tools. Future Microbiol. 5, 455–469 (2010).

2. Yeung, K. H. T., Duclos, P., Nelson, E. A. S. & Hutubessy, R. C. W. An update of the global burden of pertussis in children younger than 5 years: a modelling study. The Lancet Infectious Diseases 17, 974–980 (2017).

3. Fedele, G., Schiavoni, I., Adkins, I., Klimova, N. & Sebo, P. Invasion of Dendritic Cells, Macrophages and Neutrophils by the Bordetella Adenylate Cyclase Toxin: A Subversive Move to Fool Host Immunity. Toxins 9, 293 (2017).

4. Linhartová, I. et al. RTX proteins: a highly diverse family secreted by a common mechanism. FEMS Microbiol Rev 34, 1076–1112 (2010).

5. Chenal, A., Sotomayor-Perez, A. C. & Ladant, D. Structure and function of RTX toxins. in The Comprehensive Sourcebook of Bacterial Protein Toxins 677–718 (Elsevier, 2015). doi:10.1016/B978-0-12-800188-2.00023-9.

6. Sotomayor-Pérez, A.-C., Ladant, D. & Chenal, A. Disorder-to-Order Transition in the CyaA Toxin RTX Domain: Implications for Toxin Secretion. Toxins 7, 1–20 (2014).

7. O’Brien, D. P. et al. Structural models of intrinsically disordered and calcium-bound folded states of a protein adapted for secretion. Sci Rep 5, 14223 (2015).

8. Bumba, L. et al. Calcium-Driven Folding of RTX Domain β-Rolls Ratchets Translocation of RTX Proteins through Type I Secretion Ducts. Molecular Cell 62, 47–62 (2016).

9. O’Brien, D. P. et al. Calcium-dependent disorder-to-order transitions are central to the secretion and folding of the CyaA toxin of Bordetella pertussis, the causative agent of whooping cough. Toxicon 149, 37–44 (2018).

10. Hackett, M., Guo, L., Shabanowitz, J., Hunt, D. F. & Hewlett, E. L. Internal Lysine Palmitoylation in Adenylate Cyclase Toxin from *Bordetella pertussis*. Science 266, 433–435 (1994).

11. Basar, T. et al. The Conserved Lysine 860 in the Additional Fatty-acylation Site of Bordetella pertussis Adenylate Cyclase Is Crucial for Toxin Function Independently of Its Acylation Status. Journal of Biological Chemistry 274, 10777–10783 (1999).

12. Barry, E. M. et al. Bordetella pertussis adenylate cyclase toxin and hemolytic activities require a second gene, cyaC, for activation. J Bacteriol 173, 720–726 (1991).

13. Knapp, O. & Benz, R. Membrane Activity and Channel Formation of the Adenylate Cyclase Toxin (CyaA) of Bordetella pertussis in Lipid Bilayer Membranes. Toxins 12, 169 (2020).

14. Hewlett, E. L., Urban, M. A., Manclark, C. R. & Wolff, J. Extracytoplasmic adenylate cyclase of Bordetella pertussis. Proc. Natl. Acad. Sci. U.S.A. 73, 1926–1930 (1976).

15. Davi, M., Sadi, M., Pitard, I., Chenal, A. & Ladant, D. A Robust and Sensitive Spectrophotometric Assay for the Enzymatic Activity of Bacterial Adenylate Cyclase Toxins. Toxins 14, 691 (2022).

16. Subrini, O. et al. Characterization of a Membrane-active Peptide from the Bordetella pertussis CyaA Toxin. Journal of Biological Chemistry 288, 32585–32598 (2013).

17. Voegele, A., Subrini, O., Sapay, N., Ladant, D. & Chenal, A. Membrane-Active Properties of an Amphitropic Peptide from the CyaA Toxin Translocation Region. Toxins 9, 369 (2017).

18. Karst, J. C. et al. Identification of a Region That Assists Membrane Insertion and Translocation of the Catalytic Domain of Bordetella pertussis CyaA Toxin. Journal of Biological Chemistry 287, 9200–9212 (2012).

19. Voegele, A. et al. A High-Affinity Calmodulin-Binding Site in the CyaA Toxin Translocation Domain is Essential for Invasion of Eukaryotic Cells. Advanced Science 8, 2003630 (2021).

20. Abettan, A., Nguyen, M.-H., Ladant, D., Monticelli, L. & Chenal, A. CyaA translocation across eukaryotic cell membranes. Front. Mol. Biosci. 11, (2024).

21. Voegele, A. et al. Translocation and calmodulin-activation of the adenylate cyclase toxin (CyaA) of*Bordetella pertussis*. Pathogens and Disease 76, (2018).

22. O’Brien, D. P. et al. Calmodulin fishing with a structurally disordered bait triggers CyaA catalysis. PLoS Biol 15, e2004486 (2017).

23. El-Azami-El-Idrissi, M. et al. Interaction of Bordetella pertussis Adenylate Cyclase with CD11b/CD18. Journal of Biological Chemistry 278, 38514–38521 (2003).

24. O’Brien, D. P. et al. Post-translational acylation controls the folding and functions of the CyaA RTX toxin. FASEB J 33, fj201802442RR (2019).

25. Carbonetti, N. H., Artamonova, G. V., Andreasen, C. & Bushar, N. Pertussis Toxin and Adenylate Cyclase Toxin Provide a One-Two Punch for Establishment of *Bordetella pertussis* Infection of the Respiratory Tract. Infect Immun 73, 2698–2703 (2005).

26. Skopova, K. et al. Cyclic AMP-Elevating Capacity of Adenylate Cyclase Toxin-Hemolysin Is Sufficient for Lung Infection but Not for Full Virulence of Bordetella pertussis. Infect Immun 85, (2017).

27. Guo, Q. et al. Structural basis for the interaction of Bordetella pertussis adenylyl cyclase toxin with calmodulin. EMBO J 24, 3190–201 (2005).

28. Goldsmith, J. A., DiVenere, A. M., Maynard, J. A. & McLellan, J. S. Structural basis for non- canonical integrin engagement by Bordetella adenylate cyclase toxin. Cell Reports 40, 111196 (2022).

29. Sukova, A., et al. Negative charge of the AC-to-Hly linking segment modulates calcium-dependent membrane activities of Bordetella adenylate cyclase toxin. Bba-Biomembranes 1862, (2020).

30. Karst, J. C. et al. Calcium, Acylation, and Molecular Confinement Favor Folding of Bordetella pertussis Adenylate Cyclase CyaA Toxin into a Monomeric and Cytotoxic Form. J Biol Chem 289, 30702–16 (2014).

31. Cannella, S. E. et al. Stability, structural and functional properties of a monomeric, calcium- loaded adenylate cyclase toxin, CyaA, from Bordetella pertussis. Sci Rep 7, 42065 (2017).

32. Deruelle, V. et al. Interplay between T3SS effectors, ExoY activation, and cGMP signaling in*Pseudomonas aeruginosa*infection. Preprint at 10.1101/2025.03.26.645418 (2025).

33. Jumper, J. et al. Highly accurate protein structure prediction with AlphaFold. Nature 596, 583– 589 (2021).

34. Cossio, P. & Hummer, G. Bayesian analysis of individual electron microscopy images: Towards structures of dynamic and heterogeneous biomolecular assemblies. Journal of Structural Biology 184, 427–437 (2013).

35. Vinogradova, O. et al. A Structural Mechanism of Integrin αIIbβ3 “Inside-Out” Activation as Regulated by Its Cytoplasmic Face. Cell 110, 587–597 (2002).

36. Lau, T.-L., Kim, C., Ginsberg, M. H. & Ulmer, T. S. The structure of the integrin αIIbβ3 transmembrane complex explains integrin transmembrane signalling. EMBO J 28, 1351–1361 (2009).

37. Shang, S., Liu, J. & Hua, F. Protein acylation: mechanisms, biological functions and therapeutic targets. Sig Transduct Target Ther 7, (2022).

38. Liu, Y.-Q., Yang, Q. & He, G.-W. Post-translational acylation of proteins in cardiac hypertrophy. Nat Rev Cardiol (2025) doi:10.1038/s41569-025-01150-1.

39. S. Mesquita, F., et al. Mechanisms and functions of protein S-acylation. Nat Rev Mol Cell Biol 25, 488–509 (2024).

## Bibliography

40. Karst, J. C. et al. Calcium, Acylation, and Molecular Confinement Favor Folding of Bordetella pertussis Adenylate Cyclase CyaA Toxin into a Monomeric and Cytotoxic Form. J Biol Chem 289, 30702–16 (2014).

41. O’Brien, D. P. et al. SEC-SAXS and HDX-MS: A powerful combination. The case of the calcium-binding domain of a bacterial toxin. Biotechnol Appl Biochem 65, 62–68 (2018).

42. Trewhella, J. et al. 2017 publication guidelines for structural modelling of small-angle scattering data from biomolecules in solution: an update. Acta Crystallogr D Struct Biol 73, 710–728 (2017).

43. David, G. & Pérez, J. Combined sampler robot and high-performance liquid chromatography: a fully automated system for biological small-angle X-ray scattering experiments at the Synchrotron SOLEIL SWING beamline. J. Appl. Cryst. 42, 892–900. (2009).

44. Konarev, P. V., Volkov, V. V., Sokolova, A. V., Koch, M. H. J. & Svergun, D. I. PRIMUS: a Windows PC-based system for small-angle scattering data analysis. J Appl Crystallogr 36, 1277–1282 (2003).

45. Hopkins, J. B., Gillilan, R. E. & Skou, S. *BioXTAS RAW* : improvements to a free open-source program for small-angle X-ray scattering data reduction and analysis. J Appl Crystallogr 50, 1545–1553 (2017).

46. Guinier, A. La diffraction des rayons X aux très petits angles : application à l’étude de phénomènes ultramicroscopiques. Ann. Phys. 11, 161–237 (1939).

47. Brookes, E., Vachette, P., Rocco, M. & Perez, J. US-SOMO HPLC-SAXS module: dealing with capillary fouling and extraction of pure component patterns from poorly resolved SEC-SAXS data. J Appl Crystallogr 49, 1827–1841 (2016).

48. Brookes, E. & Rocco, M. Recent advances in the UltraScan SOlution MOdeller (US-SOMO) hydrodynamic and small-angle scattering data analysis and simulation suite. Eur Biophys J 47, 855–864 (2018).

49. Punjani, A., Rubinstein, J. L., Fleet, D. J. & Brubaker, M. A. cryoSPARC: algorithms for rapid unsupervised cryo-EM structure determination. Nat Methods 14, 290–296 (2017).

50. Jumper, J. et al. Highly accurate protein structure prediction with AlphaFold. Nature 596, 583– 589 (2021).

51. Goldsmith, J. A., DiVenere, A. M., Maynard, J. A. & McLellan, J. S. Structural basis for antibody binding to adenylate cyclase toxin reveals RTX linkers as neutralization-sensitive epitopes. PLoS Pathog 17, e1009920 (2021).

52. Motlova, L., Klimova, N., Fiser, R., Sebo, P. & Bumba, L. Continuous Assembly of β-Roll Structures Is Implicated in the Type I-Dependent Secretion of Large Repeat-in-Toxins (RTX) Proteins. Journal of Molecular Biology 432, 5696–5710 (2020).

53. Einspahr, H. & Bugg, C. E. The geometry of calcium carboxylate interactions in crystalline complexes. Acta Crystallogr B Struct Sci 37, 1044–1052 (1981).

54. Huang, J. & MacKerell, A. D. CHARMM36 all-atom additive protein force field: Validation based on comparison to NMR data. J. Comput. Chem. 34, 2135–2145 (2013).

55. Jo, S., Kim, T., Iyer, V. G. & Im, W. CHARMM-GUI: A web-based graphical user interface for CHARMM. J Comput Chem 29, 1859–1865 (2008).

56. Brooks, B. R. et al. CHARMM: The biomolecular simulation program. J Comput Chem 30, 1545–1614 (2009).

57. Lee, J. et al. CHARMM-GUI Input Generator for NAMD, GROMACS, AMBER, OpenMM, and CHARMM/OpenMM Simulations Using the CHARMM36 Additive Force Field. J. Chem. Theory Comput. 12, 405–413 (2016).

58. Abraham, M. J. et al. GROMACS: High performance molecular simulations through multi-level parallelism from laptops to supercomputers. SoftwareX 1–2, 19–25 (2015).

59. MacKerell, A. D. et al. All-Atom Empirical Potential for Molecular Modeling and Dynamics Studies of Proteins. J. Phys. Chem. B 102, 3586–3616 (1998).

60. Essmann, U. et al. A smooth particle mesh Ewald method. The Journal of Chemical Physics 103, 8577–8593 (1995).

61. Nam, K., Gao, J. & York, D. M. An Efficient Linear-Scaling Ewald Method for Long-Range Electrostatic Interactions in Combined QM/MM Calculations. J. Chem. Theory Comput. 1, 2–13 (2005).

62. Hess, B., H. Bekker, H.J.C. Berendsen, J.G.E.M. Fraaije. LINCS: A Linear Constraint Solver for Molecular Simulations. J Comput Chem 1431–1563 (1997).

63. Parrinello, M. & Rahman, A. Polymorphic transitions in single crystals: A new molecular dynamics method. Journal of Applied Physics 52, 7182–7190 (1981).

64. Souza, P. C. T. et al. Martini 3: a general purpose force field for coarse-grained molecular dynamics. Nat Methods 18, 382–388 (2021).

65. Marrink, S. J., Risselada, H. J., Yefimov, S., Tieleman, D. P. & De Vries, A. H. The MARTINI Force Field: Coarse Grained Model for Biomolecular Simulations. J. Phys. Chem. B 111, 7812–7824 (2007).

66. Monticelli, L. et al. The MARTINI Coarse-Grained Force Field: Extension to Proteins. J. Chem. Theory Comput. 4, 819–834 (2008).

67. Kroon, P., et al. Martinize2 and Vermouth: Unified Framework for Topology Generation. Preprint at 10.7554/elife.90627.2 (2024).

68. Berendsen, H. J. C., Postma, J. P. M., van Gunsteren, W. F., DiNola, A. & Haak, J. R. Molecular dynamics with coupling to an external bath. The Journal of Chemical Physics 81, 3684–3690 (1984).

69. Bussi, G., Donadio, D. & Parrinello, M. Canonical sampling through velocity rescaling. The Journal of Chemical Physics 126, 014101–7 (2007).

70. Cossio, P. et al. BioEM: GPU-accelerated computing of Bayesian inference of electron microscopy images. Computer Physics Communications 210, 163–171 (2017).

71. Cossio, P. & Hummer, G. Bayesian analysis of individual electron microscopy images: Towards structures of dynamic and heterogeneous biomolecular assemblies. Journal of Structural Biology 184, 427–437 (2013).

72. Laio, A. & Parrinello, M. Escaping free-energy minima. Proc. Natl. Acad. Sci. U.S.A. 99, 12562–12566 (2002).

73. Bonomi, M. & Parrinello, M. Enhanced Sampling in the Well-Tempered Ensemble. Phys. Rev. Lett. 104, (2010).

74. Grudinin, S., Garkavenko, M. & Kazennov, A. *Pepsi-SAXS* : an adaptive method for rapid and accurate computation of small-angle X-ray scattering profiles. Acta Crystallogr D Struct Biol 73, 449– 464 (2017).

75. Krivov, G. G., Shapovalov, M. V. & Dunbrack, R. L. Improved prediction of protein side- chain conformations with SCWRL4. Proteins 77, 778–95 (2009).

76. Pesce, F. & Lindorff-Larsen, K. Refining conformational ensembles of flexible proteins against small-angle x-ray scattering data. Biophysical Journal 120, 5124–5135 (2021).

77. Monet, D., Desdouits, N., Nilges, M. & Blondel, A. *mkgridXf* : Consistent Identification of Plausible Binding Sites Despite the Elusive Nature of Cavities and Grooves in Protein Dynamics. J. Chem. Inf. Model. 59, 3506–3518 (2019).

78. De Jong, D. H. et al. Improved Parameters for the Martini Coarse-Grained Protein Force Field. J. Chem. Theory Comput. 9, 687–697 (2013).

79. Wassenaar, T. A., Ingólfsson, H. I., Böckmann, R. A., Tieleman, D. P. & Marrink, S. J. Computational Lipidomics with *insane* : A Versatile Tool for Generating Custom Membranes for Molecular Simulations. J. Chem. Theory Comput. 11, 2144–2155 (2015).

80. Bussi, G., Donadio, D. & Parrinello, M. Canonical sampling through velocity rescaling. The Journal of Chemical Physics 126, 014101–7 (2007).

81. Bernetti, M. & Bussi, G. Pressure control using stochastic cell rescaling. The Journal of Chemical Physics 153, 114107 (2020).

82. Humphrey, W., Dalke, A. & Schulten, K. VMD: Visual molecular dynamics. Journal of Molecular Graphics 14, 33–38 (1996).

83. Gowers, R. et al. MDAnalysis: A Python Package for the Rapid Analysis of Molecular Dynamics Simulations. in 98–105 (Austin, Texas, 2016). doi:10.25080/Majora-629e541a-00e.

