## Supplementary Information for "Post-translational acylation drives folding and activity of the CyaA bacterial toxin"

### Supplementary Figures

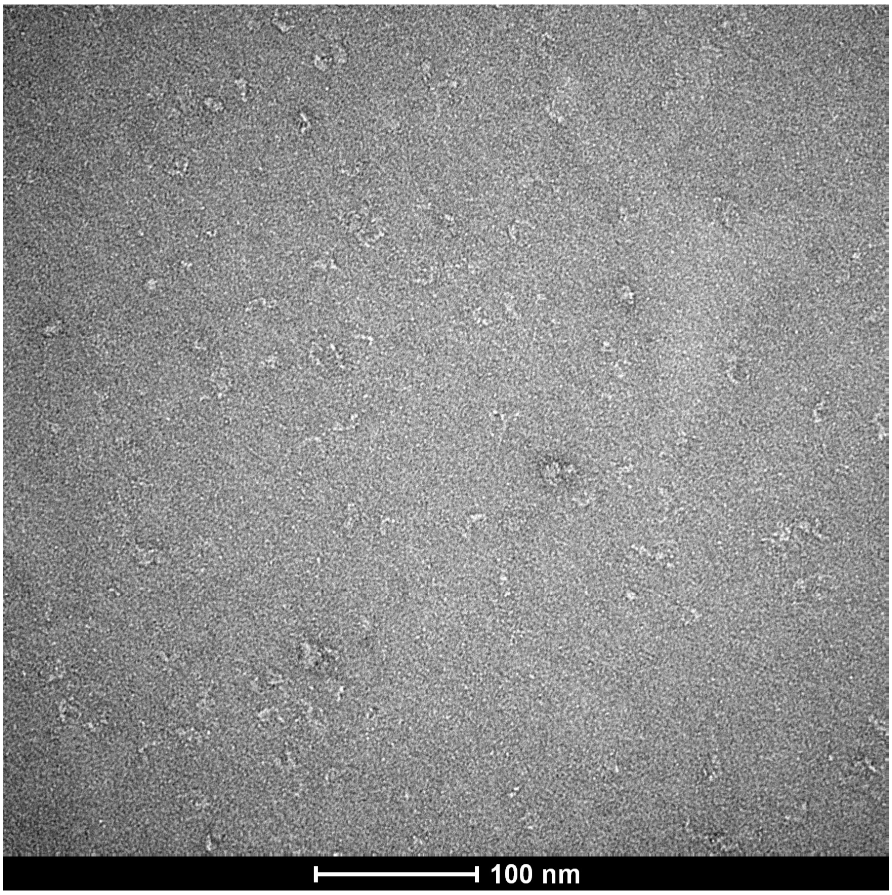

**Supplementary Fig 1. Electron micrograph of negatively stained CyaA.** The micrograph is representative of CyaA particles stained by uranyl acetate. CyaA particles have shape and size in agreement with SAXS data (**Extended Data Fig. 3**).

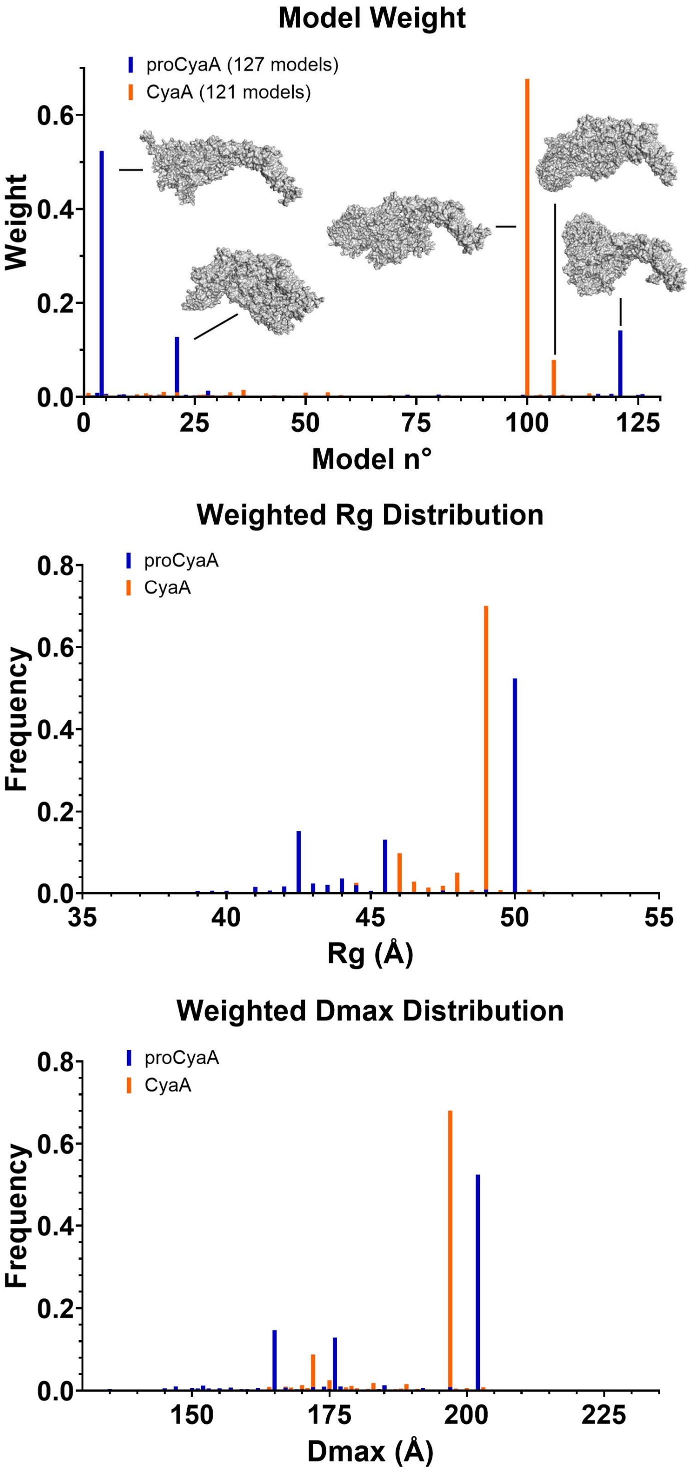

**Supplementary Fig 2. Analysis of the Hc2B integrative ensembles.** A. Model weight distribution across all the members of the proCyaA (blue bars) and CyaA (orange bars) ensembles. Representative models are visualized as grey molecular surfaces. B-C. BioEM-weighted distributions of (B) radius of gyration (Rg) and (C) maximum interatomic distance (Dmax) across all the members of the proCyaA and CyaA ensembles, colored as in panel A.

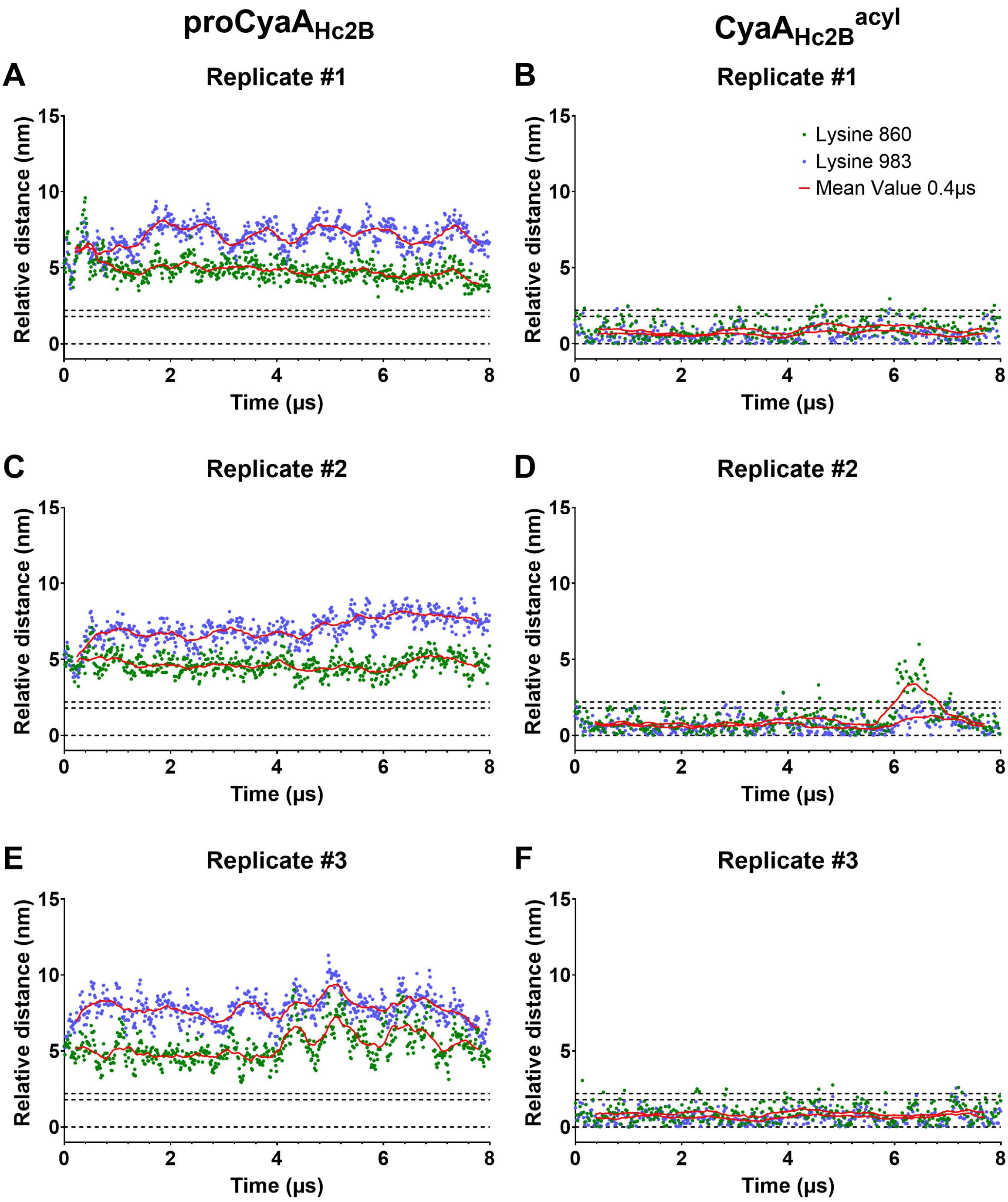

**Supplementary Fig 3. Lysine depth insertion into the lipid bilayer during coarse-grained molecular dynamic simulations. A-F.** Lysine residues K860 and K983 depth insertions into membrane for each replicate of molecular dynamic simulations of proCyaA$_{Hc2B}$ (**A, C and E**) and CyaA$_{Hc2B}^{Acyl}$ (**B, D and F**) in complex with integrins and membrane. Lysine residues K860 and K983 are in green and blue dots, respectively. Rolling means over 0.4µs are represented as red lines. See movie "Supplementary Movies 4 and 5". Horizontal dashed lines at 0, 1.5 and 2.5 nm correspond to the center of the lipid bilayer (the interface between the *cis* and *trans* lipid leaflets), the interface between the acyl chain and the lipid headgroups, and the interface between the lipid headgroups and the solvent, respectively.

### Supplementary Tables

| Method | | Protein | $R_g$ (Å) | $D_{max}$ (Å) | SC (S) | $R_H$ (Å) | [η] (mL/g) | MM (kDa) |
|---|---|---|---|---|---|---|---|---|
| SEC-SAXS[a] | GNOM[1] | CyaA | 47.3 ± 0.3 | 165 | | | | |
| | | proCyaA | 47.6 ± 0.7 | 175 | | | | |
| | Guinier[2] | CyaA | 46.5 ± 0.3 | | | | | |
| | | proCyaA | 48.7 ± 0.9 | | | | | |
| | Volume of correlation[3] | CyaA | | | | | | 182 |
| | | proCyaA | | | | | | 177 |
| AUC | | CyaA | | | 7.4 | 56 | | 176 |
| | | proCyaA | | | 7.1 | 58 | | |
| SEC-TDA | | CyaA | | | | 53 | 5.5 | 175 |

**Supplementary Table 1. Structural and hydrodynamic parameters of CyaA and proCyaA.** The parameters are derived from Size Exclusion Chromatography followed by Small-Angle X-ray Scattering (SEC-SAXS - this study), Size Exclusion Chromatography followed by a Triple Detector Array (SEC-TDA)[4,5], velocity and equilibrium Analytical UltraCentrifugation (AUC)[4,6] experimental data. Rg stands for radius of gyration (Å), Dmax for maximum distance (Å), SC for sedimentation coefficient (S), [η] for intrinsic viscosity (mL/g), MM for molecular mass (kDa, kg/mol) and (a) for *this study*.

| Method | Model | Pepsi-SAXS[7] ($Chi^2$) | | Crysol[8] | | HYDROPRO[9] | | | | |
|---|---|---|---|---|---|---|---|---|---|---|
| | | proCyaA | CyaA | $R_g$ (Å) | $D_{max}$ (Å) | $R_g$ (Å) | $D_{max}$ (Å) | S | $R_H$ (Å) | [η] (mL/g) |
| AlphaFold | Rank_1 | 2.16 | 5.68 | 57 | 211 | 56 | 215 | 6.8 | 61 | 8.8 |
| | Rank_2 | 1.96 | 5.24 | 53 | 193 | 52 | 196 | 6.9 | 60 | 8.1 |
| | Rank_3 | **1.54** | **2.07** | **53** | **202** | **51** | **206** | **6.8** | **61** | **8.8** |
| | Rank_4 | 2.17 | 6.38 | 56 | 201 | 54 | 205 | 6.6 | 63 | 9.4 |
| | Rank_5 | 2.20 | 6.59 | 56 | 197 | 55 | 201 | 6.8 | 62 | 8.8 |
| All Atom CyaA | Top cluster | | 6.97 | 55 | 204 | 54 | 208 | 5.8 | 60 | 7.5 |
| All Atom proCyaA | Top cluster | 2.34 | | 57 | 216 | 58 | 221 | 5.6 | 61 | 8.6 |
| Martini CyaA | Alone | | 2.72 | 50 | 202 | 50 | 205 | 7.2 | 59 | 7.4 |
| | + HDX-MS | | 2.22 | 47 | 182 | 48 | 186 | 7.5 | 56 | 6.4 |
| | + Bio-EM | | 1.92 | 48 | 190 | 47 | 190 | 7.2 | 56 | 6.7 |
| | Hc2B model | | 2.04 | 49 | 197 | 50 | 200 | 7.5 | 56 | 6.5 |
| Martini proCyaA | Alone | 1.69 | | 50 | 202 | 50 | 205 | 7.2 | 59 | 7.4 |
| | + HDX-MS | 1.56 | | 44 | 165 | 44 | 169 | 7.6 | 56 | 6.0 |
| | + Bio-EM | 1.39 | | 47 | 185 | 5 | 171 | 7.4 | 57 | 6.3 |
| | Hc2B model | 1.72 | | 50 | 202 | 49 | 205 | 7.1 | 59 | 7.5 |

Supplementary Table 2. **Structural parameters computed from proCyaA and CyaA models.** Rg stands for radius of gyration (Å), Dmax for maximum distance (Å), SC for sedimentation coefficient (S), [η] for intrinsic viscosity (mL/g), MM for molecular mass (kDa, kg/mol) and (a) for *this study*.

| Data collection parameters | | |
|---|---|---|
| Instrument | Beamline SWING (Synchrotron SOLEIL) | |
| Detector | EigerX4M in vacuum (Dectris) | |
| Sample to detector distance (m) | 2.00 | |
| Beam geometry | 400 µm x 200 µm | |
| Wavelength (Å) | 1.033 | |
| q-range ($Å^{-1}$) | $0.0041 < q < 0.50$ | |
| Exposure time (ms) / reading time (ms) | 990 / 10 | |
| Temperature (K) | 288 | |
| **Sample** | **proCyaA monomers** | **CyaA monomers** |
| Molar extinction coefficient ($M^{-1}.cm^{-1}$) | 143,590 | 143,590 |
| CyaA molecular mass (Da) | 177,590 | 177,590 |
| Injected volume (µL) | 50 | 50 |
| Concentration (µM \| mg/mL) | 5.3 \| 0.95 and 16.1 \| 2.86 | 16.8 \| 2.94 |
| Running buffer of the SEC | Hepes 20mM, NaCl50 mM, $CaCl_2$ 2mM, pH 7.4 | Hepes 20mM, NaCl50 mM, $CaCl_2$ 2mM, pH 7.4 |
| **Structural parameters** | | |
| I(0) Guinier ($cm^{-1}$) | 0.025 ± 0.001 | 0.024 ± 0.001 |
| $R_g$ Guinier (Å) | 48.7 ± 0.9 | 46.5 ± 0.3 |
| q-range for Guinier fit ($Å^{-1}$) | 0.008 – 0.026 | f0.016 – 0.028 |
| I(0) P(r) ($cm^{-1}$) | 0.024 ± 0.002 | 0.024 ± 0.001 |
| $R_g$ P(r) (Å) | 47.6 ± 0.6 | 47.3 ± 0.3 |
| $D_{max}$ (Å) | 170 | 165 |
| q-range for P(r) fit ($Å^{-1}$) | 0.009-0.400 | 0.016-0.399 |
| **Data reduction and analysis software** | | |
| FOXTROT[10], ATSAS 3.2.1[11], RAW 2.1.4[12], US-SOMO 4.0[13], CRYSOL 2.8.3[8] | | |

**Supplementary Table 3.** SEC-SAXS data collection and scattering derived parameters for CyaA and proCyaA.

| | Replica 1 | Replica 2 | Replica 3 | Average ± std error |
|---|---|---|---|---|
| **Palm860 - Palm983** | 4 | 7 | 46 | 19 ± 14 |
| **Palm860 in Pocket1** | 81 | 63 | 48 | 64 ± 10 |
| **Palm983 in Pocket2** | 31 | 15 | 20 | 22 ± 5 |
| **Palm983 in Pocket3** | 78 | 87 | 22 | 63 ± 21 |
| **Palm983 in Pocket2 and Pocket3** | 14 | 9 | 0 | 7 ± 7 |

Supplementary Table 4. **Population of contacts (Percent of simulation time).** Contact was measured with 6A distance cutoff, using Carbon C4 for all calculations, except for the contact between the two acyl chains where C3 and C4 contacts were used.

### Supplementary Movies

All movies were generated in Blender using Molecular Nodes add-on. [14]

**Supplementary Movie 1 - proCyaA-vs-CyaA-5us-360z+360x**
5µs Coarse-Grained Molecular Dynamic of proCyaA$_{Hc2B}$ (left) and CyaA$_{Hc2B}^{Acyl}$ (Right) in solution rotating 360° on z then on x axes. Both molecules are represented in sphere with ACD in green, TR in orange, HR in blue, AR in black and RD in dark red with acylated K860 and K983 in red and purple respectively.

**Supplementary Movie 2 - proCyaA-vs-CyaA-Acyl-5us--20+20-20**
5µs Coarse-Grained Molecular Dynamic of proCyaA$_{Hc2B}$ (left) and CyaA$_{Hc2B}^{Acyl}$ (Right) in solution rotating from -20° to +20° to -20° on the z axis. Both molecules are represented in sphere with ACD in green, TR in orange, HR in blue, AR in black and RD in dark red with acylated lysines 860 and 983 in red and purple respectively.

**Supplementary Movie 3 - CyaA-Acyl-5us-pockets**
5µs Coarse-Grained Molecular Dynamic of CyaA in solution. CyaA$_{Hc2B}^{Acyl}$ is represented in sphere grey sphere with acylated K860 and K983 in red and purple respectively and hydrophobic pockets 1, 2 and 3 in Blue, Orange and Yellow respectively.

**Supplementary Movie 4 - CyaA-complex-8us**
8µs Coarse-Grained Molecular Dynamic of CyaA$_{Hc2B}^{Acyl}$ in complex with chimeric integrins and a lipid bilayer. CyaA$_{Hc2B}^{Acyl}$ is represented in sphere with ACD in green, TR in orange, HR in blue, AR in black and RD in dark red with acylated K860 and K983 in red and purple spheres respectively. The lipid bilayer is represented by the lipid headgroup carbons in yellow sphere. Chimeric CD11b and CD18 integrins are represented in dark and light purple spheres, respectively.

**Supplementary Movie 5 - proCyaA-complex-8us**
8µs Coarse-Grained Molecular Dynamic of proCyaA$_{Hc2B}$ in complex with chimeric integrins and a lipid bilayer. proCyaA$_{Hc2B}$ is represented in sphere with ACD in green, TR in orange, HR in blue, AR in black and RD in dark red with K860 and K983 in red and purple spheres respectively. The lipid bilayer is represented by the lipid headgroup carbons in yellow sphere. Chimeric CD11b and CD18 integrins are represented in dark and light purple spheres, respectively.

**Supplementary Movie 6 - CyaA-complex-start-200ns**
First 200ns of the Coarse-Grained Molecular Dynamic of CyaA$_{Hc2B}^{Acyl}$ in complex with chimeric integrins and a lipid bilayer. CyaA$_{Hc2B}^{Acyl}$ is represented in sphere with ACD in green, TR in orange, HR in blue, AR in black and RD in dark red with acylated K860 and K983 in red and purple spheres respectively. The lipid bilayer is represented by the lipid headgroup carbons in yellow sphere. Chimeric CD11b and CD18 integrins are represented in dark and light purple spheres, respectively.

**Supplementary Movie 7 - proCyaA-complex-start-200ns**
First 200ns of Coarse-Grained Molecular Dynamic of proCyaA$_{Hc2B}$ in complex with chimeric integrins and a lipid bilayer. proCyaA$_{Hc2B}$ is represented in sphere with ACD in green, TR in orange, HR in blue, AR in black and RD in dark red with K860 and K983 in red and purple spheres respectively. The lipid bilayer is represented by the lipid headgroup carbons in yellow sphere. Chimeric CD11b and CD18 integrins are represented in dark and light purple spheres, respectively.

#### Bibliography

1. Svergun, D. I. Determination of the Regularization Parameter in Indirect -Transform Methods Using Perceptual Criteria. *J. Appl. Cryst.* **25**, 495–503 (1992).
2. Guinier, A. La diffraction des rayons X aux très petits angles; application à l'étude de phénomènes ultramicroscopiques. *Ann Phys (Paris)* **12**, 161–237 (1939).
3. Rambo, R. P. & Tainer, J. A. Accurate assessment of mass, models and resolution by small-angle scattering. *Nature* **496**, 477-+ (2013).
4. Cannella, S. E. *et al.* Stability, structural and functional properties of a monomeric, calcium-loaded adenylate cyclase toxin, CyaA, from Bordetella pertussis. *Sci Rep* **7**, 42065 (2017).
5. Karst, J. C. *et al.* Calcium, Acylation, and Molecular Confinement Favor Folding of Bordetella pertussis Adenylate Cyclase CyaA Toxin into a Monomeric and Cytotoxic Form. *J Biol Chem* **289**, 30702–16 (2014).
6. O'Brien, D. P. *et al.* Post-translational acylation controls the folding and functions of the CyaA RTX toxin. *FASEB J* **33**, fj201802442RR (2019).
7. Grudinin, S., Garkavenko, M. & Kazennov, A. *Pepsi-SAXS* : an adaptive method for rapid and accurate computation of small-angle X-ray scattering profiles. *Acta Crystallogr D Struct Biol* **73**, 449–464 (2017).
8. Svergun, D. I., Barberato, C. & Koch, M. H. J. CRYSOL - a program to evaluate X-ray solution scattering of biological macromolecules from atomic coordinates. *J. Appl. Crystallogr.* **28**, 768–773 (1995).
9. Ortega, A., Amoros, D. & Garcia de la Torre, J. Prediction of hydrodynamic and other solution properties of rigid proteins from atomic- and residue-level models. *Biophys J* **101**, 892–8 (2011).
10. Thureau, A., Roblin, P. & Pérez, J. BioSAXS on the SWING beamline at Synchrotron SOLEIL. *J Appl Crystallogr* **54**, 1698–1710 (2021).
11. Manalastas-Cantos, K. *et al. ATSAS 3.0* : expanded functionality and new tools for small-angle scattering data analysis. *J Appl Crystallogr* **54**, 343–355 (2021).
12. Hopkins, J. B., Gillilan, R. E. & Skou, S. *BioXTAS RAW* : improvements to a free open-source program for small-angle X-ray scattering data reduction and analysis. *J Appl Crystallogr* **50**, 1545–1553 (2017).
13. Brookes, E. & Rocco, M. Recent advances in the UltraScan SOlution MOdeller (US-SOMO) hydrodynamic and small-angle scattering data analysis and simulation suite. *Eur Biophys J* **47**, 855–864 (2018).
14. Johnston, B. *et al.* BradyAJohnston/MolecularNodes: v4.5.0. Zenodo https://doi.org/10.5281/zenodo.17037902 (2025).
